# The livebearers platyfish and swordtails partially regenerate their hearts with persistent scarring

**DOI:** 10.1101/2025.09.23.678041

**Authors:** Vincent Hisler, Lana Rees, Simon Blanchoud, Heidi E.L. Lischer, Rémy Bruggmann, Anna Jaźwińska

## Abstract

Heart regeneration varies among vertebrates, with zebrafish serving as a reference species for efficient cardiac restoration. How this capacity diversified across teleosts is an emerging question, particularly following the recent identification of non-regenerative cardiac repair in medaka and cavefish. Here, we investigate heart restorative capacity following cryoinjury in two livebearers, platyfish and swordtails (*Xiphophorus* species), belonging to the *Poeciliidae* family. We demonstrate that their hearts lack the vascularized compact myocardium, a ventricular layer implicated in the restorative response in zebrafish. Following cryoinjury, both poeciliids failed to rapidly deposit fibrotic tissue that normally reinforces the damaged ventricle. This deficiency correlates with pronounced wound protrusion. Although the remaining myocardium displayed an initial proliferative response, subsequently deposited collagenous scar tissue permanently sealed the ventricular wall, precluding complete regeneration. Transcriptomic analysis identified several divergently regulated pathways between cryoinjured hearts of zebrafish and platyfish, most notably in immune response regulation. These differences were validated by delayed leukocyte infiltration and sustained inflammation in platyfish, contrasting with the rapid and self-resolving inflammatory response in zebrafish. Our findings demonstrate that *Xiphophorus* species have evolved hearts with compromised regenerative capacity, characterized by initial wound protrusion and permanent scarring. These results establish that lineage-specific evolutionary traits can profoundly shape regenerative competence across teleosts, with broad implications for understanding the mechanistic basis of cardiac repair.

**Highlights:** · Viviparous poeciliids lack vascularized compact myocardium.

· Inflammation and fibrosis are delayed in the cryoinjured platyfish ventricle.

· Ventricular cryoinjury in *Xiphophorus* leads to transient bulging-type deformation.

· Failure to form a myocardial bridge results in permanent scarring.

## INTRODUCTION

Efficient healing of heart injury is crucial for maintaining blood flow and ensuring survival. In most adult mammals, including humans, myocardial infarction damages ventricular muscle, which is ultimately replaced with a collagenous scar ^1,2^. By contrast, certain fish and amphibians, notably zebrafish and axolotls, can regenerate a fully functional myocardium without scarring, as demonstrated in partial ventricular resection models ^3–8^. There is therefore debate about why this regenerative capacity was lost during vertebrate evolution ^9–12^. Interpretations of these findings require caution, however, since teleosts and tetrapods represent distinct evolutionary lineages, and the zebrafish heart may not faithfully represent the ancestral condition ^13^.

Teleosts form a monophyletic taxon with a 300-million-year history, whose remarkable radiation has created over 30,000 extant species ^14–17^. Interestingly, this extensive diversification led to the establishment of distinct heart architectures, as already noted by McWilliam in 1885: *"The cardiac structure and blood-supply appear to present considerable diversity in different fishes. Thus, in the salmon, the dense outer part of the ventricular wall is of great thickness, comprising several longitudinal, circular and oblique layers…; in the cod’s heart there appears to be no distinct outer layer of dense muscular tissue … And there are no coronary vessels to be discerned upon the surface as in the salmon, eel and others."* ^18,19^. Given that heart architecture and systemic factors are considered to influence restorative strategies ^20–22^, this note suggests that comparative studies across multiple fish lineages are essential before broad conclusions about cardiac regeneration can be drawn. Indeed, teleost cardiac diversity may map directly onto a corresponding diversity of regenerative responses. This situation contrasts with the mammalian heart, whose gross anatomy is remarkably conserved from mouse to whale ^13,23^.

The zebrafish has become the leading animal model for studying efficient heart regeneration ^10,22,24–26^. After cryoinjury, the damaged myocardium is substantially replaced within 30 days, although a full recovery may require additional months ^27–29^. Strikingly, the regenerative program can be restarted multiple times in the same individual ^30^. Beside the myocardium, also the endocardium, epicardium, vasculature, peripheral nerves, and immune system actively participate in the restorative process ^31–36^. Remarkably, the activation of the full regenerative program is not universal for all teleost, as medaka and a cave-dwelling variant of the Mexican tetra, which are species from distinct teleost orders, heal cardiac wounds through fibrosis ^37–39^. The relationship between heart type and regenerative capacity therefore remains poorly defined, because only a small number of fish species have been examined after cardiac injury. A systematic comparative approach could illuminate how this health-relevant trait evolved within the teleost clade ^40–42^.

This study aims to assess cardiac regeneration in viviparous fishes from the family *Poeciliidae* of the order *Cyprinodontiformes*. We selected two members of the *Xiphophorus* genus, the platyfish (*X. maculatus*) and the swordtail (*X. hellerii*), for several reasons. The order *Cypriniformes* (comprising zebrafish) and *Cyprinodontiformes* (platyfish and swordtails) diverged approximately 250 million years ago, making these groups well-suited for deep evolutionary comparisons ^14,15,43,44^ (**Figure 1A**). *Xiphophorus* has attracted scientific interest for a century thanks to several peculiarities, including variation in pigmentation, susceptibility to melanomas, internal fertilization, sexual trait elaboration, geographical adaptability, evolutionary hybridization and radiation ^45–50^. The platyfish genome has been annotated, rendering it suitable for transcriptomic analysis ^51^. Our laboratory has recently used this species to study fin regeneration, and reported the presence of a few epichordally-derived principal rays in the caudal fin, which can be recognized as a rule-breaking trait among teleosts ^52,53^. Despite this suite of interesting features, no *Cyprinodontiformes* or live-bearing fish has previously been examined for cardiac regenerative capacity.

**Figure 1.**
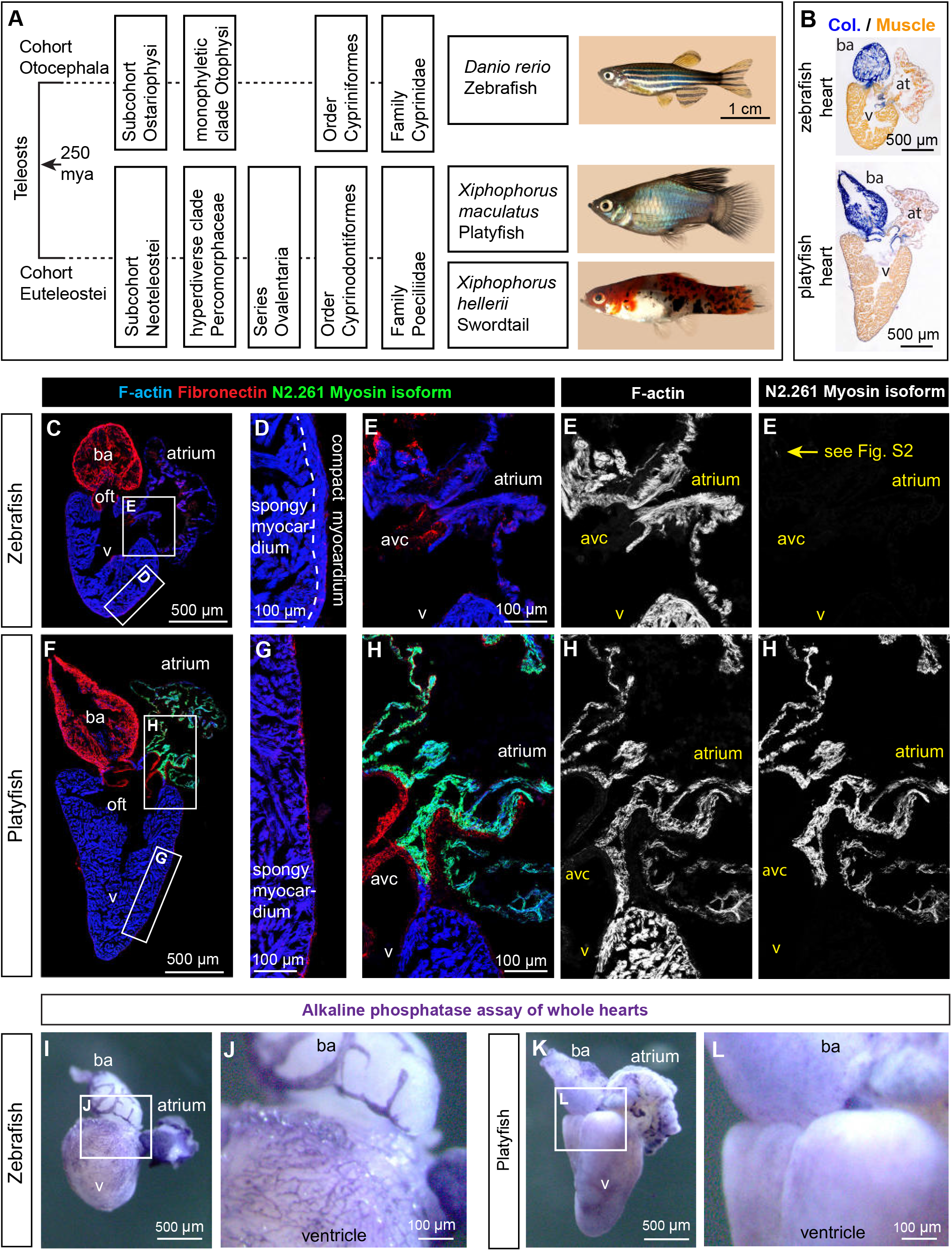
Platyfish possess a heart lacking vascularized compact myocardium. **(A)** Simplified phylogenetic lineages of the fish species examined in this study, based on ref. ^14^. Teleosts diversified into two main branches approximately 250 million years ago (mya). The *Otophysi* clade encompasses more than one-third of all fish species, including zebrafish and tetras ^43^. *Percomorphaceae* is a hyperdiverse clade described as “*the bush at the top*” of the fish phylogenetic tree ^116^, comprising approximately 55% of extant teleost diversity, including perciforms, cichlids, and poeciliids^117^. **(B)** AFOG staining of longitudinal sections of platyfish and zebrafish hearts. Collagen (Col.) is stained in blue. **(C-H)** Fluorescence staining of hearts shown in (B). The myocardium is labeled by F-actin staining (fluorescent phalloidin). Note the presence of the compact myocardium in zebrafish (D), and its absence in the platyfish (G). The N2.261 antibody recognizes a specific myosin isoform that is nearly absent in the uninjured zebrafish heart (C-E), with the exception of a few fibers at the outflow tract (oft). In platyfish, N2.261 immunolabels the atrium, suggesting evolutionary divergence of myosin isoform composition between species. Fibronectin, an extracellular matrix protein, is detected in the bulbus arteriosus. **(I-L)** Alkaline phosphatase activity staining reveals a dense coronary vasculature in zebrafish hearts, which is absent in platyfish. Frames depict the magnified areas shown on adjacent panels, labeled with the corresponding letter. Abbreviations: avc, atrioventricular canal; at, atrium; ba, bulbus arteriosus; oft, outflow tract; v, ventricle. These abbreviations and labeling conventions apply throughout all figures.

Here, we demonstrate the differences in the architecture and molecular components of the heart between zebrafish and *Xiphophorus sp.* and compare the restorative response after cryoinjury using immunofluorescence and transcriptomic analyses. Our findings indicate that livebearers can partially regenerate the heart but with persistent scarring, and we propose that this intermediate restorative strategy reflects evolutionary innovations in both ventricular structure and immune function. Expanding the taxonomic breadth of regenerative biology promises to deepen our mechanistic understanding of how cardiac regeneration evolved across vertebrates.

## RESULTS

### *Xiphophorus* lacks vascularized compact myocardium in the ventricle

To establish baseline anatomical differences that might influence regenerative capacity, we compared heart morphology between adult zebrafish and platyfish. In teleosts, the heart consists of a single atrium and ventricle, which pumps blood to an elastic outflow chamber, the bulbus arteriosus. AFOG staining of longitudinal sections labeled the bulbus arteriosus and valves with a collagen-binding blue dye, while the ventricle and atrium appeared in beige (**Figure 1B**). The platyfish ventricle was longer and more pyramidal than that of zebrafish, with a distinctly pointed apex. Swordtails displayed an identical architecture (**suppl. Figure S1A**), indicating that these ventricular differences are consistent across *Xiphophorus* species.

To assess molecular composition, sections were labeled with three markers: fluorescent phalloidin to visualize the F-actin-rich contractile tissue, anti-Fibronectin for connective tissue, and the N2.261 (embCMHC) antibody against a specific isoform of slow myosin (**Figure 1C-H**). In zebrafish, N2.261 has previously been identified as a marker of immature cardiomyocytes (CMs) in larvae and in regenerating myocardium ^54,55^. In the intact heart of adult zebrafish, only individual CMs were labeled near the outflow tract **(Figure 1C, E; suppl. Figure S2A-C**). Platyfish and swordtails, by contrast, displayed N2.261 immunoreactivity throughout the entire atrium (**Figure 1F, H; suppl. Figure S1B-C; suppl. Figure S2D-F**). This pattern was already present in platyfish embryos, whereas immature cardiomyocytes are not labeled by this antibody.(**suppl. Figure S2H, I**). Together, these findings indicate that N2.261 recognizes distinct myosin heavy chain isoforms in zebrafish and *Xiphophorus*: an embryonic ventricular isoform in the former and an atrial isoform in the latter. The differential labeling pattern between species is more likely attributable to evolutionary divergence in myosin heavy chain sequences than to differences in cardiomyocyte maturation state, reflecting the substantial lineage-specific reshaping of cardiac myosin repertoires that has occurred between cyprinids and poeciliids.

Most critically, zebrafish and *Xiphophorus* differed in the architecture of the outer ventricular layer: zebrafish possessed a vascularized compact myocardium, as previously described ^56^, whereas this layer was entirely absent in *Xiphophorus* (**Figure 1D, G; suppl. Figure S1C; suppl. Figure S2G**). To determine whether compensatory structural differences exist in the tissue surrounding the ventricular chamber, we performed picrosirius red (PSR) staining and imaged it under polarized light to detect fibrillar collagen ^57,58^ (Figure 2). Both groups showed fibrillar collagen in the cardiac valves and in the thin sheath surrounding the bulbus arteriosus (Figure 2E-G, M-O, suppl. Figure S1H-J). At the outer ventricular wall, however, the two groups diverged sharply: zebrafish ventricular myocardium was essentially devoid of fibrillar collagen, whereas the equivalent layer in *Xiphophorus* was covered by a distinct collagen-rich coat (Figure 2H, P, suppl. Figure 1K). This suggests that the epicardium of poeciliids is substantially enriched in fibrillar extracellular matrix relative to that of zebrafish.

**Figure 2.**
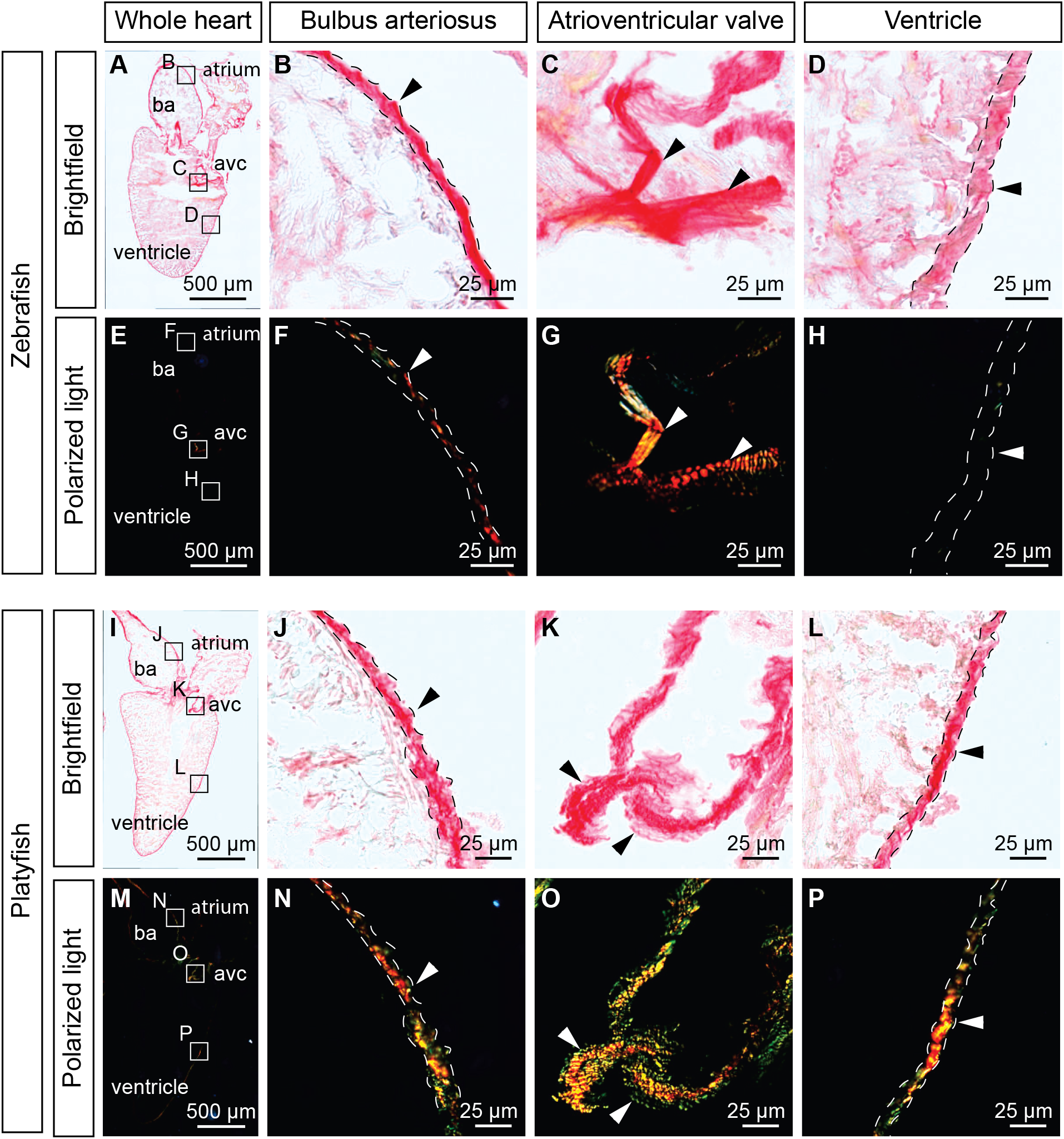
The outer layer of *Xiphophorus* ventricle consists of a thin, collagen-rich matrix. Picrosirius Red (PSR) staining of longitudinal heart sections from zebrafish **(A–H)** and platyfish **(I–P)**. Sections were imaged under bright-field **(A–D, I–L)** and polarized light **(E–H, M–P)**. For each species, higher magnification views of the outer layer of the bulbus arteriosus **(B, F, J, N)**, valves **(C, G, K, O)**, and the outer layer of the myocardium **(D, H, L, P)** are shown. Under polarized light, collagen fibers display a range of color from green to red depending on fiber thickness and extracellular matrix composition ^118^. In contrast to zebrafish (**H**), platyfish (**P**) exhibit a distinct fibrillar collagenous coat surrounding the ventricle.

To examine vascularization, whole hearts were stained for alkaline phosphatase activity ^39^. The zebrafish ventricle was covered by an extensive vascular network, whereas platyfish and swordtail showed no comparable vasculature (**Figure 1I-L, suppl. Figure 1M, L**). Podocalyxin-2 (Podxl2) immunostaining, which detects endothelial cell apical surfaces ^59^, confirmed the absence of ventricular vascularization in platyfish (**suppl. Figure S3**). Taken together, these results demonstrate that *Xiphophorus* and zebrafish hearts differ fundamentally at both morphological and molecular levels, with *Xiphophorus* lacking the compact myocardium and associated coronary vasculature that participate in cardiac regeneration in zebrafish.

### Cryoinjured ventricles of *Xiphophorus* regenerate with deformation and persistent scarring

Cryoinjury creates reproducible damage through controlled freezing and thawing using a precooled probe ^60^ (**Figure 3A; suppl. Fig S4A**). Examination of whole platyfish hearts at 7 days post-cryoinjury (7 dpci) revealed profound Phalloidin-negative tissue protruding from the ventricular wall, indicating extensive myocardial loss (**Figure 3B**). To evaluate regenerative dynamics, transverse sections of cryoinjured hearts were analyzed using AFOG staining across multiple time points. In zebrafish, transient collagenous tissue appeared between 7 and 14 dpci, detected by Aniline blue staining, as previously reported ^27,28,61^ (**Figure 3C**). In *Xiphophorus* species, by contrast, the damaged area contained Fuchsin red-stained protein deposits at these time points, with minimal collagen. This pattern suggests significantly delayed wound clearance in *Xiphophorus* compared to zebrafish.

**Figure 3.**
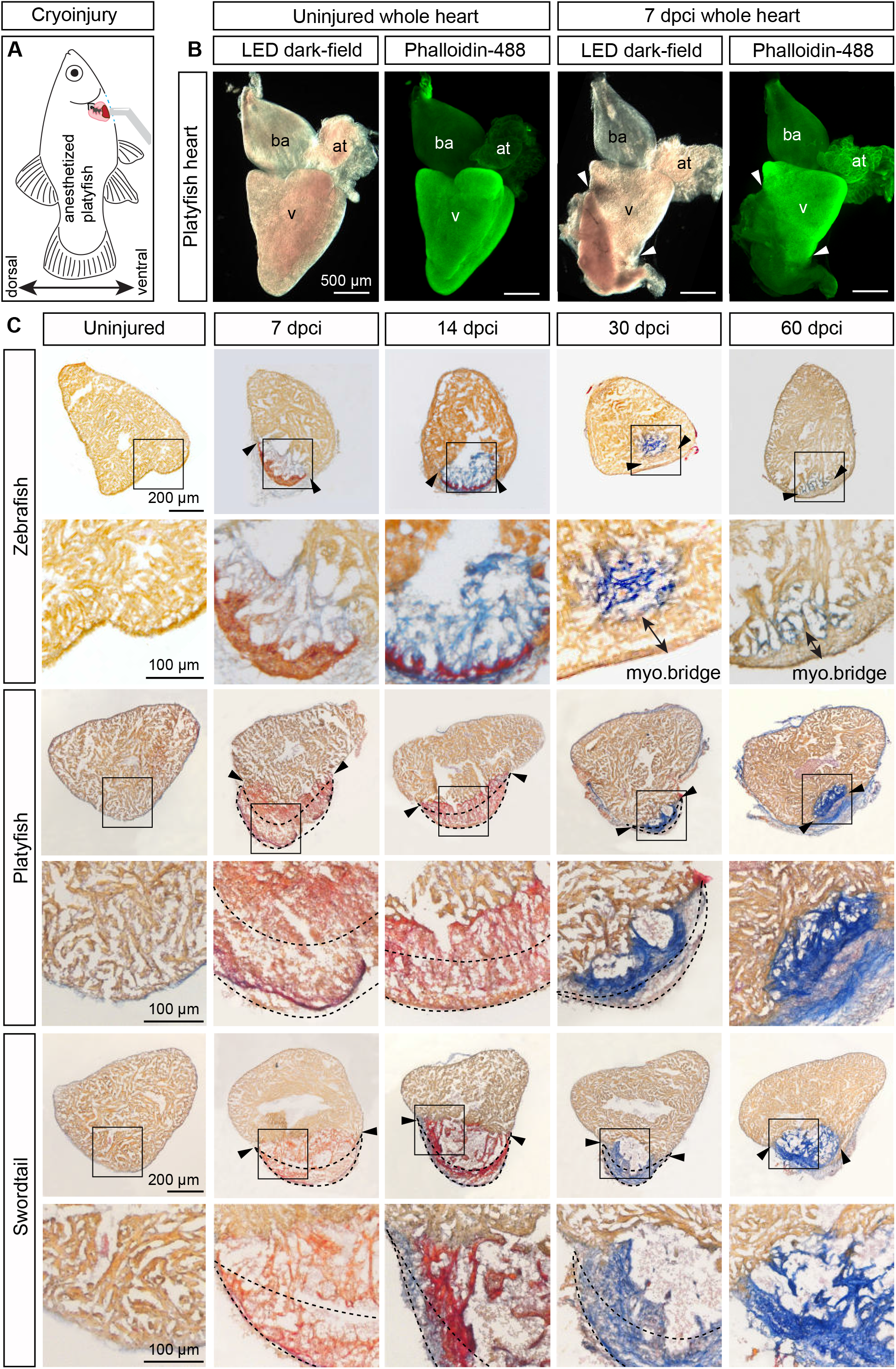
Cryoinjured ventricles in *Xiphophorus* fish display transient wound bulging and permanent scarring. **(A)** Schematics of heart cryoinjury procedure in platyfish. **(B)** Representative images of hearts from uninjured fish and at 7 days post-cryoinjury (dpci), stained with fluorescent phalloidin (green). The damaged area is identified by a weak fluorescence signal reflecting the loss of contractile cardiomyocytes. Arrowheads indicate the border between the intact myocardium and the wound. Abbreviations: at, atrium; ba, bulbus arteriosus; v, ventricle. **(C)** AFOG staining of transverse ventricular sections from zebrafish, platyfish, and swordtail, collected at indicated time points after cryoinjury. Intact myocardium (orange); fibrin and other protein deposits (red); collagen (blue). Arrowheads indicate the wound edge; double-headed arrows mark the myocardial (myo) bridge; dashed lines encircle wound tissue that has expanded beyond the normal ventricular circumference, indicative of the wound bulging phenotype.

Strikingly, *Xiphophorus* wounds displayed marked bulging beyond the presumptive ventricular margin at 7 and 14 dpci (**Figure 3B, C dashed line**), a phenotype absent in zebrafish and other non-regenerative species (medaka and cavefish) ^37–39^. Unlike these species, *Xiphophorus* showed collagen deposition only after 14 dpci, indicating a substantial delay in fibrotic tissue formation (**Figure 3C**). Together, these findings reveal that collagen-deficient wounds are more prone to mechanical deformation in *Xiphophorus*, uncovering a novel variation in teleost cardiac healing strategies.

To determine the persistence of this phenotype, we examined hearts at 30, 60 and 90 dpci. In zebrafish, the wound was substantially resorbed and the "myocardial bridge" structure spanned the wounded myocardium, as previously described ^62,63^ (**Figure 3C, Figure S4B**). In livebearers, by contrast, a belt-like collagen layer sealed the wound in a pattern strikingly distinct from the fine fibrillar network characteristic of zebrafish regeneration (**Figure 3C, suppl. Figure S4B**). Despite this persistent scarring, the wound size at 30 dpci appeared smaller than that at 7 dpci, indicating partial regeneration. Despite this persistent scarring, the wound area at 30 dpci appeared reduced compared to 7 dpci, indicating that partial myocardial restoration had occurred.

To quantify deformation frequency, phenotypes were categorized into three groups: little to no wound, typical non-protruding wound, and protruding swollen wound (**Figure 4A**). At 7 dpci, approximately 45% of platyfish and and 75% swordtail hearts displayed protruding swollen wounds. At 14 dpci, this proportion decreased by nearly half, and by 30-90 dpci, wound swelling was rarely observed, indicating gradual resorption of the bulging deformity. Nevertheless, persistent fibrosis was evident in most hearts, confirming limited regeneration. Notably, this restoration proceeded in the absence of detectable N2.261 immunoreactivity in the peri-injury zone (**suppl. Figure S4C**), contrasting with the robust upregulation of this embryonic myosin heavy chain isoform observed during zebrafish cardiac regeneration^54,55^. The absence of this dedifferentiation marker in the livebearer border zone suggests that partial myocardial restoration in these species proceeds through mechanisms distinct from those operating in zebrafish.

**Figure 4.**
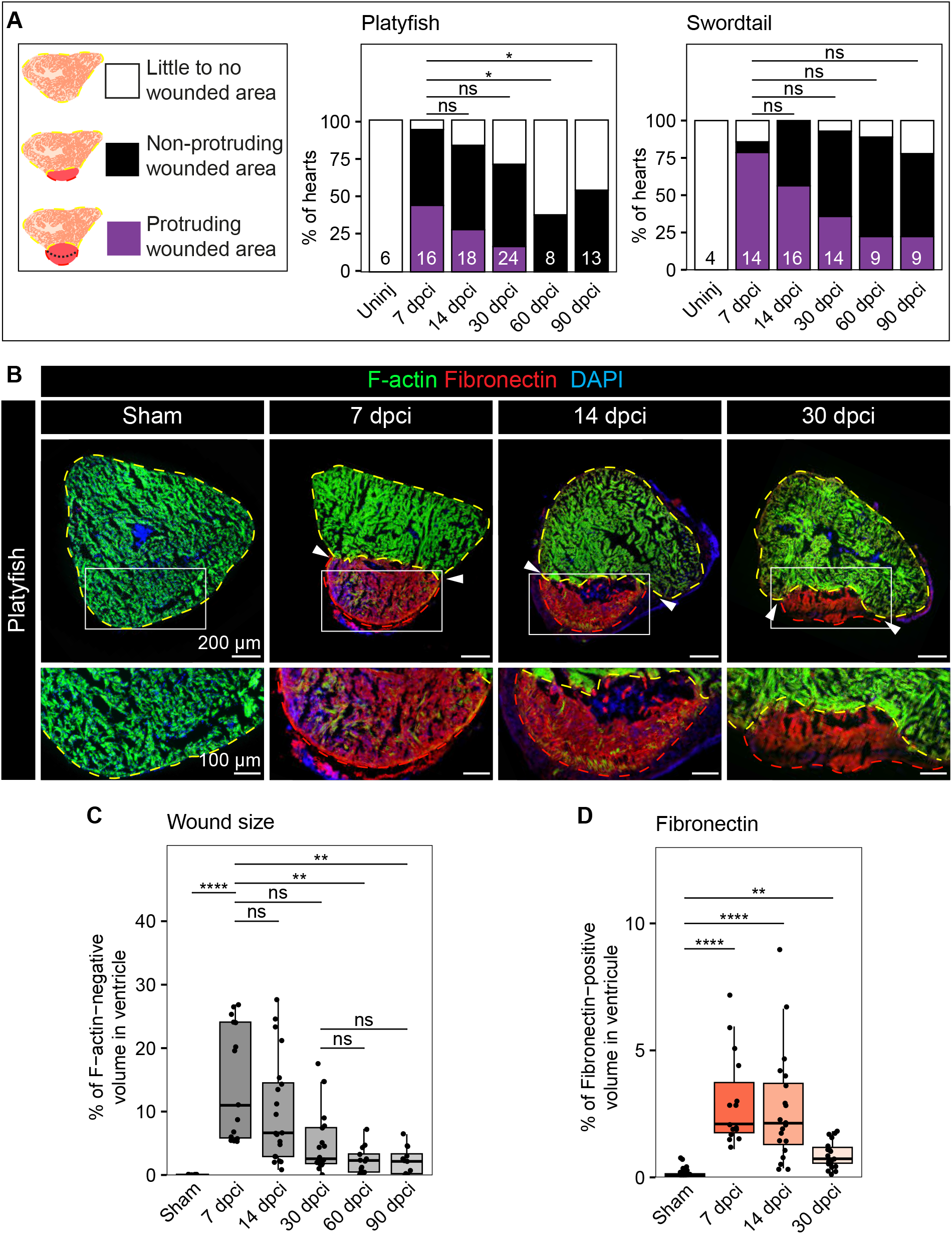
Partial restoration of the heart in platyfish and swordtail after cryoinjury. **(A)** Classification of injury phenotypes based on AFOG staining, representative examples of which are shown in Figure 3. Stacked bar plots display the percentage of each category at indicated time points after cryoinjury in platyfish and swordtail. Numbers at the base of each bar indicate the number of biological replicates (fish). Pearson’s chi-squared test with Holm’s post hoc correction: ns, not significant; *, p < 0.05. **(B)** Fluorescence staining of transverse platyfish heart sections at indicated time points after cryoinjury. The fibronectin-positive wound area (red) contrasts with the intact myocardium labeled by F-actin (green). Sham-operated ventricles at 30 days post-thoracotomy are shown as controls. **(C-D)** Quantification of the wound size and fibronectin deposition, as represented in (B). Statistical comparisons were performed using the Kruskal-Wallis test followed by Dunn’s test with Holm’s post hoc correction. Adjusted p-value: * < 0.05, ** < 0.01, *** < 0.001, **** < 0.0001. Sample sizes: n = 24 (sham), 15 (7 dpci), 20 (14 dpci), 20 (30 dpci), 13 (60 dpci), and 12 (90 dpci).

To quantify healing outcomes, platyfish hearts were stained with fluorescent Phalloidin and anti-Fibronectin antibody, and tissue volumes were calculated from stained areas across serial heart sections. Consistent with AFOG findings, Phalloidin staining confirmed the absence of myocardial bridge formation in livebearers **(Figure 4B, suppl. Figure S5A)**. . Wound volume, defined as the Phalloidin-negative ventricular fraction, comprised 15% (SD ±10%) of total ventricle at 7 dpci, declining to 10% (±10%) at 14 dpci, 5% (±6%) at 30 dpci, and 2% (±2%) at 60-90 dpci (**Figure 4C**). Swordtails showed similar dynamics (**suppl. Figure S5B, C**). Fibronectin quantification revealed a progressive decrease in connective tissue paralleling wound volume reduction (**Figure 4D**). Collectively, these data demonstrate that while Xiphophorus species achieve substantial wound size reduction over time, this occurs through fibrotic sealing rather than true myocardial regeneration, as evidenced by persistent collagenous scarring and the absence of regenerative myocardial bridge formation.

### Cryoinjured ventricles show transcriptomic differences between zebrafish and platyfish

To identify molecular differences underlying their distinct regenerative responses, we performed bulk transcriptomic analysis of ventricles at 7 dpci alongside uninjured controls. In each species, we identified genes with differential transcript abundance (**Figure 5A, B; suppl. Table S1, S2**). Interspecies comparison revealed that following cryoinjury, 199 orthologous gene transcripts showed a higher log₂ fold change (log₂FC) in zebrafish than in platyfish, while 268 showed the opposite pattern, demonstrating that the two species mount distinct transcriptional responses to cardiac injury (**Figure 5C**).

**Figure 5.**
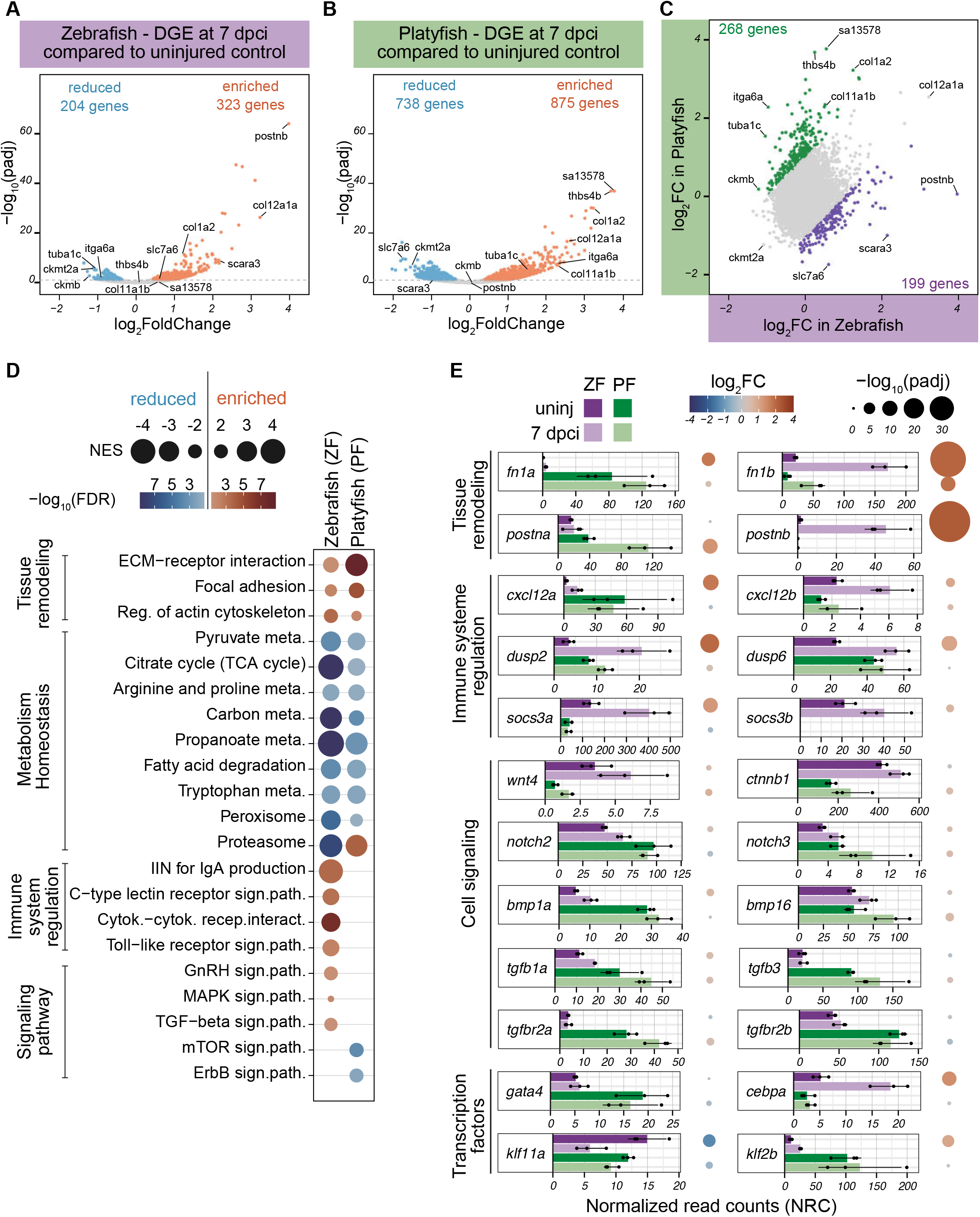
Bulk RNA sequencing reveals divergent responses to cryoinjury between zebrafish and platyfish. (A-B) Volcano plots of transcriptomes from cryoinjured zebrafish (A) and platyfish (B) ventricles at 7 dpci, compared to uninjured controls. The log₂ fold change (log₂FC) represents the ratio of transcript abundance in cryoinjured versus uninjured conditions. Genes with a significant change in transcript levels (adjusted p-value, padj < 0.05) are highlighted in blue (decreased) or orange (increased). **(C)** Scatter plot comparing log₂FC values between the two species. Genes with a higher log₂FC in platyfish than in zebrafish are shown in green; those with a higher log₂FC in zebrafish are shown in purple. Highlighted genes show at least a twofold difference in transcript abundance after cryoinjury between species. **(D)** Comparison of gene set enrichment analyses (GSEA) between species. Color intensity reflects the statistical significance of gene set reduction (blue) or enrichment (orange) at 7 dpci relative to uninjured controls. Dot size corresponds to the normalized enrichment score (NES) for each gene set. Selected gene sets are organized into broader functional categories. **(E)** Transcript abundance of orthologous genes across conditions (uninjured, dark shade; cryoinjured, light shade) and species (zebrafish, ZF, purple; platyfish, PF, green). Each point represents the DESeq2-based TPM-like normalized read count (see Methods) for one biological replicate. The log₂FC and −log₁₀(padj) values correspond to those shown in (A) and (B). Genes are grouped into broader functional categories.

Gene Set Enrichment Analysis (GSEA) revealed that tissue remodeling factors were enriched whereas metabolic regulators were reduced in both species **(Figure 5D; suppl. Table 3)**. This suggests a broadly conserved response to disrupted cardiac homeostasis following injury ^26,64^. Notably, species-specific differences emerged within immune-related pathways: C-type lectin receptor, cytokine and Toll-like receptor signaling were enriched exclusively in zebrafish. Likewise, GnRH, MAPK, and TGFβ signaling pathways were enriched in zebrafish but not in platyfish, whereas mTOR and ErbB signaling pathways were specifically reduced in platyfish.

To better understand these species-specific responses, we examined individual gene families in detail. Comparing normalized read counts (NRC; see Materials and Methods) and log₂FC, we found that paralogues within the same gene family often showed divergent patterns of differential transcript abundance between species. For example, *fibronectin 1a (fn1a)* transcripts were more abundant in platyfish, whereas *fn1b* predominated in zebrafish (Figure 5E). Similar patterns were observed for the periostin family (*postna/postnb*). These results highlight that species-specific paralogue usage should be carefully considered when interpreting cross-species transcriptome comparisons.

Given our observation of delayed collagen deposition in the platyfish wounded heart at 7 dpci (**Figure 3C**), we next examined extracellular matrix (ECM)-associated genes (**suppl. Figure S6**). Only modest interspecies differences were detected: zebrafish hearts appeared more enriched in collagen V and VI as well as tenascin, while platyfish hearts showed higher levels of elastin, metalloproteinases, and disintegrins. These differences in ventricular ECM composition may contribute to the reduced regenerative capacity of platyfish. Nevertheless, cryoinjury induced broadly similar changes in transcript abundance for most ECM components examined across both species, including fibronectin, periostin, *col1a1b/2*, *col12a1a/1b*, and *mmp2* (**Figure 5E, suppl. Figure S6A, E**).

Finally, we analyzed genes and biological processes more directly associated with heart regeneration. Consistent with our GSEA results, transcripts of the chemokine ligand *cxcl12a* were enriched in zebrafish at 7 dpci, while the orthologous transcripts remained unchanged in platyfish (**Figure 5E**) ^65^. Similarly, suppressors of cytokine signalling (*socs3a*/*b*) and the dual-specificity protein phosphatase family member *dusp2* ^66^ were more abundant in zebrafish cryoinjured hearts than in their platyfish counterparts. Collectively, these findings suggest a more robust immune response in zebrafish following cardiac injury. Examination of genes associated with canonical signaling pathways (Wnt, Notch, and TGFβ) ^67–69^ revealed no striking interspecies differences. Among transcription factors, *cebpa* and *klf2b* were significantly enriched exclusively in zebrafish following cryoinjury. These factors represent candidate molecular determinants of the divergent regenerative responses observed between the two species.

### Delayed and persistent infiltration of leukocytes in the injured platyfish myocardium

GSEA revealed distinct enrichment of immune-related gene sets, including Toll-like receptor, Neuregulin/ErbB, and cytokine signaling pathways **(Figure 5D)**. Given that these pathways have been previously linked to cardiac regeneration and immune cell activation ^70–73^, we next assessed the transcript abundance of leukocyte-specific markers in both species. Despite baseline differences in gene expression between the two species, pan-leukocyte markers, including *cxcr3.2*, *ltb4r*, *ncf2* and *ptpn6,* were more strongly enriched following cryoinjury in zebrafish than in platyfish **(Figure 6A)**. This pattern was consistent across genes associated with inflammation (NF-κB/TNF-α pathway, TLR signalling, and an inflammatory regulator *nr4a1*), neutrophils (*apoeb, havcr1, irf8, mrc1b)* and macrophages (*mpx* and *rac2)* **(suppl. Figure S7A-C)**. Taken together, these results indicate a blunted immune cell activation response in platyfish relative to zebrafish.

**Figure 6.**
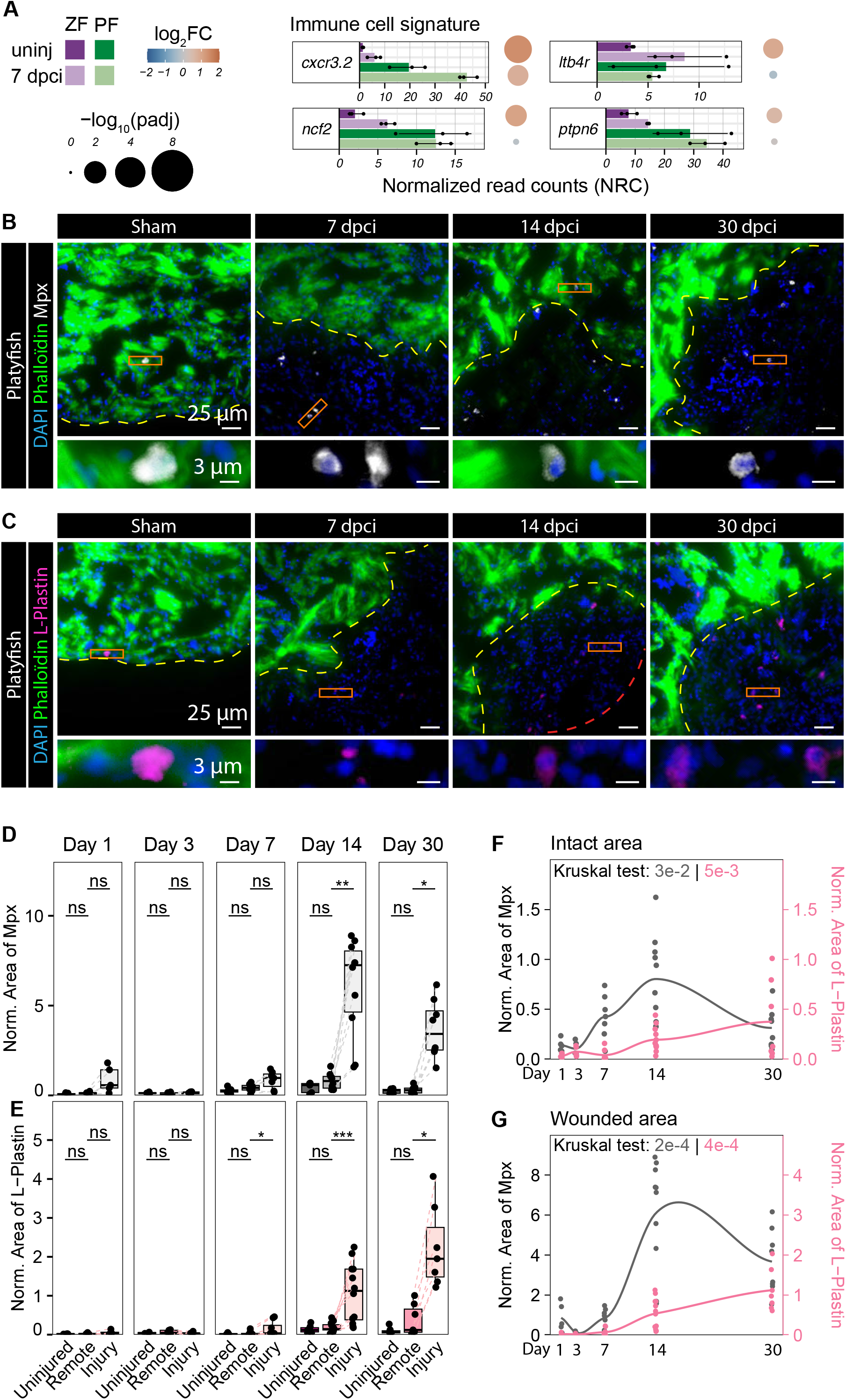
Delayed recruitment of immune cells during platyfish heart repair. **(A)** Transcript abundance of immune cell activation-related genes across conditions and species. For details, see the legend to Figure 5 and Methods. **(B-C)** Immunofluorescence staining for Mpx-positive neutrophils (B) and L-plastin-positive leukocytes (C) at 7, 14, and 30 dpci. Sections were counterstained with DAPI and phalloidin-488. Control sections were obtained from uninjured ventricles of sham-operated fish at 30 days post-thoracotomy (dpt). **(D-E)** Quantification of Mpx-positive area (D) and L-plastin-positive area (E), corresponding to representative images shown in (B) and (C). In the cryoinjured ventricles (CI), quantifications were assessed separately in the intact myocardium (CI: Intact) and the wounded area (CI: Wound). Statistical comparisons of Sham vs. CI: Intact were performed using unpaired Wilcoxon tests, as these groups comprised different animals, whereas comparisons between the intact myocardium and wound area within the same cryoinjured heart were performed using paired Wilcoxon tests. Holm’s post-hoc correction was applied for multiple comparisons. Adjusted p-value: * P< 0.05, ** P< 0.01. **(F-G)** Recruitment kinetics of neutrophils (grey, left Y-axis) and L-plastin-positive leukocytes (purple, right Y-axis) in the intact myocardium (F) and wound area (G) at 1, 3, 7, 14, and 30 dpci. Sham values pooled across all time points, represent the baseline level of each marker in uninjured myocardium. Kruskal-Wallis tests were used to compare marker levels across time points.

To characterize leukocyte infiltration spatiotemporally, we performed immunofluorescence on platyfish cardiac sections using antibodies against Myeloperoxidase (Mpx/Mpo) as a neutrophil marker and L-plastin as a pan-leukocyte marker ^74–76^ (**Figure 6B-C, suppl. Figure S7D-F**). Between 1 and 3 dpci, both Mpx-positive and L-plastin-positive cell numbers remained low, comparable to uninjured controls. Leukocyte numbers began to rise in the injury zone by 7 dpci and were significantly elevated relative to the remote myocardium from 14 dpci onward, persisting through 30 dpci (**Figure 6D-E**). In contrast, leukocyte counts in the remote myocardium remained consistently low across all time points, with only a modest increase observed in a subset of hearts.

Temporal profiling of neutrophil and pan-leukocyte dynamics revealed divergent kinetics between the two populations (**Figure 6F-G**). In both injured and intact myocardium, neutrophil numbers increased from 3 dpci, peaked at 14 dpci, and subsequently declined. L-plastin-positive leukocytes, however, began accumulating later from 7 dpci and remained elevated through 30 dpci. This protracted inflammatory response contrasts sharply with zebrafish, in which leukocytes are largely cleared by 14 dpci ^64,77–79^, demonstrating that platyfish display a markedly delayed resolution of cardiac inflammation.

### Cardiomyocyte proliferation is transient and spatially unrestricted in platyfish

Zebrafish cardiac regeneration relies on the dedifferentiation and proliferation of CMs within the border zone myocardium, a peri-injury region extending approximately 100 μm from the injury margin **(Figure 7A)** ^5,6,54^. To examine CM-specific responses in both species, we filtered our bulk transcriptomic analysis for genes associated with these processes.

**Figure 7.**
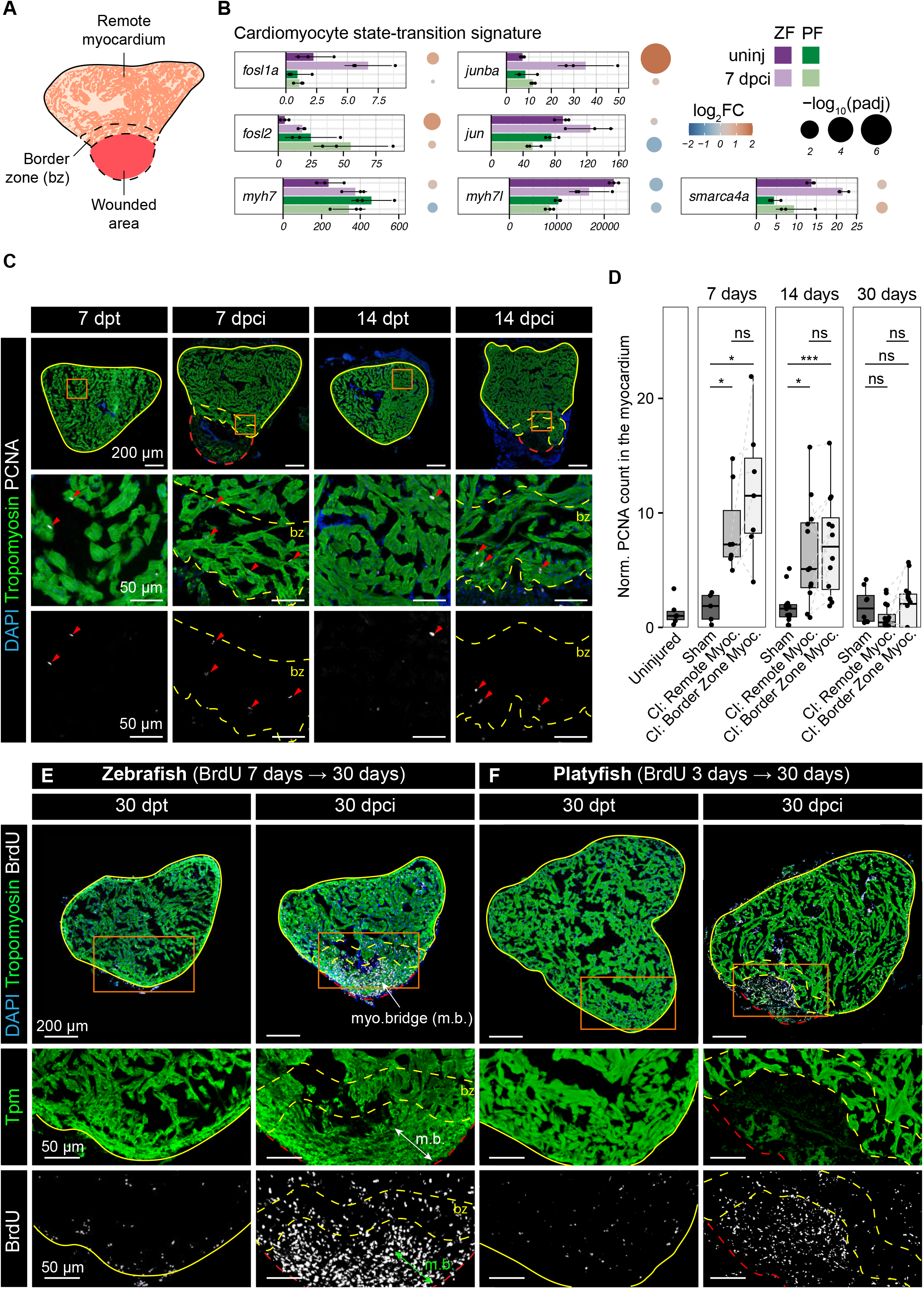
Ventricular cryoinjury triggers transient cardiomyocyte activation. **(A)** Schematic illustration of the three analyzed areas: the tropomyosin-negative wounded area, the border zone (bz) myocardium within 100 µm from the injury border, and the remote myocardium distant from the border zone. **(B)** Transcript abundance of orthologous genes defined as markers of cardiomyocyte fate change associated with cardiac regeneration. For further details, see the legend to Figure 5 and Methods. **(C)** PCNA immunolocalization in sham-operated (7 dpt and 14 dpt) and cryoinjured (7 dpci and 14 dpci) platyfish ventricles. Orange frames in the top panels indicate the regions magnified in the middle panels. Bottom panels show the corresponding PCNA channel alone. Red arrowheads indicate PCNA-positive nuclei within the myocardium. bz, border zone. **(D)** Quantification of proliferating cells in the Tropomyosin-positive myocardium across time points and cardiac regions. Sham versus CI: Remote myocardium comparisons were performed using unpaired Wilcoxon tests. CI: Remote myocardium versus CI: Border zone comparisons were performed using paired Wilcoxon tests, as measurements were obtained from the same hearts. Holm’s post hoc correction was applied for multiple comparisons. Adjusted p-value: * < 0.05, ** < 0.01, *** < 0.001, **** < 0.0001. **(E-F)** BrdU incorporation in zebrafish (E) and platyfish (F) following continuous labeling from 7 to 30 dpci (23 days) and 3 to 30 dpci (27 days), respectively. Sections were counterstained with DAPI and anti-tropomyosin antibody. Orange frames in the top panels indicate the regions magnified in the middle panels. Bottom panels display the corresponding BrdU channel alone. In zebrafish, the initial wound area is delineated based on myocardial morphology and the high density of BrdU-positive nuclei. The myocardial bridge (m.b., double-headed arrow) is observed in zebrafish but not in platyfish. For panels (C, E, F), dashed red lines encircle the wound area and dashed yellow lines demarcate the border zone.

As expected in zebrafish, cryoinjury induced significant enrichment of key regulators of transcriptional reprogramming required for CM cell cycle re-entry and dedifferentiation, including members of the Activator Protein-1 (AP-1) complex (*jun* and *fos*) and *smarca4a*/*brg1* from the SWI/SNF chromatin remodeling complex ^80–82^ (**Figure 7B**). In contrast, these enrichments were markedly attenuated or absent in platyfish. A similar divergence was observed for the ventricular myosin heavy chain, *myh7*, one isoform of which is recognized by the N2.261 antibody ^83,84^ (**Figure 7B**), and for the *connexin 43* (*cx43*), also implicated in CM proliferation ^85^ (**suppl. Figure S8B**). Interestingly, *myh7l*, which is expressed in adult ventricle ^86^, was downregulated in both species (**Figure 7B**). Consistent with ongoing CM redifferentiation at 7 dpci in zebrafish, we also detected enrichment of the transcriptional coactivator *cited4a*, which regulates the balance between CM proliferation and maturation ^87^, as well as Rbfox family splicing regulators involved in sarcomere assembly ^88–90^ (**suppl. Figure S8A**). These genes were again not significantly enriched in platyfish. Collectively, these findings indicate that platyfish exhibit a markedly reduced, though not entirely absent, activation of the CM fate-change programs required for effective cardiac regeneration.

Paradoxically, the platyfish transcriptome showed stronger enrichment of proliferation-associated genes (*pcna, cdk4, cdk2*) and DNA replication stress markers (*atm, atr, rpa*) ^91^ compared with zebrafish (**suppl. Figure S8B**). To assess whether this transcriptional signature reflects CM proliferation, we performed PCNA immunofluorescence combined with Tropomyosin (TPM) and DAPI staining, followed by quantification across the remote myocardium, border zone, and injured area **(Figure 7A)**. Sham-operated controls showed no significant increase PCNA staining, compared to uninjured hearts **(Figure 7C, D; suppl. Figure S8C, D)**. At 7 dpci, PCNA-positive nuclei were significantly elevated in both the remote myocardium and the wounded area, but declined by 14 dpci and returned to baseline by 30 dpci **(suppl. Figure S8D)**. Analysis of PCNA/TPM-double positive nuclei showed the same temporal trend, with high enrichment at 7 dpci followed by a gradual decrease **(Figure 7D)**. Strikingly, PCNA/TPM-positive nuclei in the border zone reached levels comparable to those in the remote myocardium, indicating that cryoinjury triggered transient CM cell cycle re-entry throughout the entire ventricle rather than being spatially restricted to the border zone as in zebrafish ^55,62^.

To track DNA-replicating over extended periods, we adapted the BrdU labeling approach established in zebrafish, in which treatment from 7 to 30 dpci labels approximately 60% of CMs in regenerated tissue ^54^ **(Figure 7E)**. In platyfish, BrdU administration began at day 3, with hearts collected at 7, 14, and 30 dpci **(suppl. Figure S8E)**. A clear peak of DNA synthesis in CMs was detected at 7 dpci and was equally prominent in the remote myocardium and border zone, confirming the absence of spatial restriction to the peri-injury region **(suppl. Figure S8F, G)**. At 14 and 30 dpci, BrdU incorporation in CMs dropped sharply in both regions **(Figure 7F, suppl. Figure S8F, G)**, demonstrating that the CM proliferative response is confined to the early post-injury phase, a conclusion fully consistent with the PCNA data.

Critically, unlike zebrafish, neither the wound area nor the border zone of cryoinjured platyfish hearts displayed substantial BrdU labeling at later time points **(Figure 7F)**. Together, the BrdU and PCNA analyses demonstrate that while CMs do transiently re-enter the cell cycle in platyfish, this response is neither sustained nor spatially focused, two hallmarks of efficient regeneration in zebrafish. The failure to maintain a proliferative CM population during advanced regeneration phases likely represents a key cellular mechanism underlying the incomplete cardiac regenerative response in platyfish.

## DISCUSSION

Sampling diverse phylogenetic lineages can provide new insights into the distribution of cardiac regeneration among teleosts ^92,93^. This study revealed that platyfish and swordtails, which represent live-bearing poeciliids, have evolved a ventricle that lacks vascularized compact myocardium, a structure present in the zebrafish ^56^. Zebrafish and *Xiphophorus* therefore possess fundamentally distinct cardiac architectures, with important consequences for regenerative capacity.

In platyfish and swordtail hearts, the reparative phase within the first two weeks after cryoinjury proceeded without collagen deposition and was accompanied by protrusion beyond the chamber circumference. This observation indicates that the fibrotic matrix provides essential mechanical support for wounds within the blood-pumping ventricle, consistent with findings in zebrafish ^94,95^. A comparable bulging phenomenon has been reported in zebrafish treated with inhibitors of TGF-β signaling, which prevented deposition of collagen and tenascin C, extracellular matrix components typically abundant in normally regenerating wounds ^69^. In platyfish, TGF-β signaling appears to be less active shortly after injury, and collagen and tenascin C were not enriched at the wound site, potentially explaining the molecular basis of the bulging phenotype. Further investigation is needed to clarify how specific extracellular matrix proteins contribute to the structural integrity and healing of the damaged ventricle.

During the advanced phase, between two and four weeks after cryoinjury, fibrosis filled the damaged area more densely than typically observed in zebrafish. These dynamics correlated with elevated cardiomyocyte (CM) proliferation primarily during the initial phase, prior to scar deposition. Although wound size decreased by 30 days post-injury, platyfish failed to form the myocardial bridge that normally reconnects the interrupted ventricle in regenerating species ^96^. Instead, a fibrotic scar sealed the wound margin. This failure may be a direct consequence of the absence of compact myocardium and its associated intramyocardial connective tissue, which likely provides the structural scaffold required for myocardial bridging. Taken together, *Xiphophorus* species display compromised myocardial regeneration accompanied by partial scarring, rather than true cardiac regeneration.

Transcriptomic and immunofluorescence analyses of platyfish ventricles revealed a delayed and persistent inflammatory response, characterized by prolonged retention of leukocytes and neutrophils in injured tissue. In zebrafish, macrophages contribute directly to scar formation through cell-autonomous collagen deposition ^77^. Consequently, poor initial immune cell infiltration in platyfish may explain the delayed collagenous matrix deposition in the wound, observed here. This finding is consistent with the absence of coronary vasculature, which in zebrafish facilitates immune cell circulation and delivers inductive factors that promote heart regeneration ^31,33,97–100^. The avascular heart of platyfish thus appears less compatible with efficient regeneration, even if tissue remodeling can reduce overall wound size over time. Whether the suboptimal immune response in platyfish is solely responsible for the lack of sustained CM proliferation, or whether additional intrinsic factors contribute, remains an open question. Addressing this will require transcriptomic profiling across multiple time points following injury, which would allow the temporal dynamics of immune activation, matrix remodeling, and CM fate changes to be resolved with greater precision. Such a longitudinal analysis represents a natural extension of the present study and a priority for future work.

Broader studies across fish species reinforce the view that inflammation and systemic physiology profoundly shape cardiac regenerative capacity ^37,38^. In medaka, an inappropriate immune response prevents heart regeneration, an effect that can be rescued by stimulating Toll-like receptor (TLR) signaling ^39^. Strikingly, pathway enrichment analysis in platyfish similarly showed no induction of TLR signaling, suggesting that shared immune deficiencies may underlie regenerative failure in both species, which lack vascularized compact myocardium. However, the presence of a compact layer is not sufficient to guarantee regeneration: cave-dwelling populations of the Mexican tetra retain this structure yet have lost regenerative competence, likely through metabolic adaptations to a dark and nutrient-limited habitat ^101^. This highlights that systemic and environmental factors can override structural prerequisites for regeneration. By contrast, caudal fin regeneration appears more robust to such influences and is broadly conserved across diverse teleosts, including medaka, cavefish, and *Xiphophorus* ^53,93,102–105^. This divergence may reflect the greater evolutionary conservation of the caudal fin bauplan relative to ventricular architecture across teleosts.

What selective advantage might an avascular ventricle lacking compact myocardium confer, given that the vascularized heart is considered ancestral in fish evolution and the compact layer is prominent in high-performance athletes, such as tuna ^106,107^? One plausible benefit relates to cardiovascular health: dependence on coronary vessels introduces the risk of vascular pathology. Migratory salmonids, for example, exhibit coronary lesions that may contribute to ischemic states and premature aging ^108^. The consequences of such lesions on ischemic states may correlate with premature aging. Such risks are entirely absent in the avascular *Xiphophorus* heart, suggesting a potential reduction in predisposition to coronary disease, a trade-off that may be advantageous in ecological contexts where sustained athletic performance is less critical.

Finally, in humans, myocardial infarction can lead to life-threatening changes in ventricular geometry, including pseudoaneurysm formation, the mechanistic basis of which remains incompletely understood ^109,110^. The cryoinjured *Xiphophorus* heart, which similarly exhibits ventricular deformations, may therefore represent a tractable and biomedically relevant model for investigating the cardiac pathophysiology, an avenue that warrants dedicated future study.

## MATERIALS AND METHODS

### Resources

Phylogenetic information was based on the updated classification of bony fishes, inferred using molecular and genomic data ^14^. The number of species in taxa was taken from FishBase (https://www.catalogueoflife.org). The references for chemical reagents, antibodies, and computer software are listed in the **Supplementary Materials**.

### Animal strains

*Xiphophorus hellerii* (swordtail) and *Xiphophorus maculatus* (platyfish) at approximately 3.5 cm standard length were purchased from a commercial aquarium fish vendor (Aqualand, Renens/Lausanne, Switzerland). Wild-type zebrafish were from the AB strain, bred in our fish facility. Fish were of random sex and aged between 4 months and 1 year. Fish housing animal procedures were approved by the cantonal veterinary office of Fribourg. All assays were performed using different animals randomly assigned to experimental groups. The exact sample size (n) is described for each experiment on the graphs or in figure legends.

### Ventricular cryoinjury procedure

Fish were first immersed in an analgesic solution of 5 mg/L lidocaine for 45 min, followed by transfer to an anesthetic buffered solution of 0.6 mM tricaine (MS-222) for a few minutes, until loss of responsiveness was confirmed. Depth of anesthesia was verified prior to each procedure by assessing the absence of response to tactile stimulation. Ventricular cryoinjuries were performed according to our established video protocol ^60^. Briefly, anesthetized fish were positioned dorsal side down on a moistened sponge under a stereomicroscope, exposing the ventral surface. A midline incision was made through the thoracic skin to access the pericardial cavity, and a stainless steel cryoprobe pre-cooled in liquid nitrogen was applied directly to the ventricular surface for 20–23 s. Freezing was terminated by rinsing the ventricle with room-temperature water, after which the cryoprobe was removed. Fish were immediately returned to system water and monitored continuously until spontaneous opercular movement and swimming behavior resumed, then observed for an additional several hours post-procedure. Sham-operated controls underwent thoracotomy alone: identical skin incision and pericardial exposure but without cryoprobe application, to account for surgical trauma independent of cryoinjury.

### Heart collection and fixation

Fish were euthanized by immersion in 300 mg/L buffered tricaine (MS-222) until cessation of opercular movement and loss of response to tactile stimulation were confirmed. Hearts were collected as described in our established protocol ^111^. The ventral side was reopened, and the heart was extracted. Excised hearts were briefly rinsed in phosphate-buffered saline (PBS), then fixed in 2% paraformaldehyde (PFA) overnight at 4°C. Following fixation, hearts were washed in PBS and cryoprotected by equilibration in 30% sucrose at 4°C for a minimum of 24 h. Hearts were subsequently embedded in tissue freezing medium (OCT) and stored at −80°C until sectioning.

Cryosections were cut at 12 µm for all applications, with the exception of anti-Podocalyxin immunostaining, for which 50 µm sections were prepared to preserve deeper tissue planes. Sections were collected on Superfrost Plus slides, air-dried for approximately 1 h at room temperature, and stored at −20°C in sealed boxes until further use.

### BrdU treatment

To label proliferating cells, fish were exposed to 5-bromo-2’-deoxyuridine (BrdU) for the indicated time periods. Groups of 4–5 fish were maintained in 1 L of system water supplemented with BrdU at a final concentration of 50 mg/L. Treatment water was renewed every 2 days using freshly prepared stock solution.

For each water change, a 5 mg/mL BrdU stock solution was prepared by dissolving the appropriate amount of BrdU powder in demineralized water under gentle agitation at 37°C until fully dissolved. The stock was then diluted 100-fold in system water and thoroughly mixed before reintroducing the fish.

### Immunofluorescence analysis

Cryosections were processed for immunofluorescence using a standard indirect protocol. Slides were first permeabilized with 0.3% Triton X-100 in PBS for 10 min at room temperature, then blocked with blocking buffer (5% normal goat serum, 0.3% Triton X-100 in PBS) for 1–2 h at room temperature to minimize non-specific antibody binding.

Sections were incubated overnight at 4°C with primary antibodies diluted in blocking buffer. Following three wash steps with 0.3% Triton X-100/PBS, sections were incubated with fluorophore-conjugated secondary antibodies diluted in blocking buffer for 1–2 h at room temperature, then washed again under the same conditions. Nuclei were counterstained with DAPI. In experiments requiring visualization of muscle architecture, an additional 1 h incubation with Phalloidin-CruzFluor-488 diluted in blocking buffer was performed at room temperature, followed by a final wash step. All primary and secondary antibodies, along with their working dilutions, are listed in the **Supplementary Materials**. Slides were mounted using a custom glycerol-based mounting medium and stored at 4°C until imaging.

For BrdU and PCNA immunostaining, antigen retrieval was performed between the permeabilization and blocking steps. For BrdU detection, slides were incubated in 2 N HCl/0.3% Triton X-100/PBS for 45 min at room temperature to denature DNA and expose incorporated BrdU epitopes, followed by two neutralization washes in PBS. For PCNA detection, heat-induced epitope retrieval was performed using pre-warmed sodium citrate buffer (10 mM sodium citrate, 0.05% Tween 20, pH 6.0) in a pressure cooker at maximum pressure for 3 min, then allowed to cool to room temperature before proceeding to the blocking step.

### AFOG staining

Aniline blue, acid fuchsin, and orange-G (AFOG) staining was performed as described ^69,112^. Tissue sections were first fixed in 10% formalin for 15 min at room temperature, followed by a 10-min wash in 0.3% Triton X-100 in PBS to enhance permeability. Sections were then incubated in pre-warmed Bouin’s fixative for 2.5 hours at 56°C, then allowed to continue fixation for an additional hour at room temperature. Excess picric acid was removed by washing under running tap water until the yellow color dissipated.

Sections were subsequently stained with AFOG solution, prepared by dissolving 3 g acid fuchsin, 2 g orange-G, and 1 g aniline blue in 200 mL of acidified distilled water adjusted to pH 1.1. Following staining, sections were rinsed with distilled water and dehydrated through a graded ethanol series (70%, 95%, and 100%), cleared in xylene, and coverslipped using Entellan mounting medium (Merck). With this staining protocol, fibrin and protein deposits appear red, collagen fibers appear blue, and muscle appears orange. Brightfield images were acquired using a DM6B microscope (Leica Microsystems).

### Picrosirius Red (PSR) staining

Sections were first air-dried for 30 minutes, then immersed in xylene for 30 min at room temperature, followed by rehydration through a graded ethanol series (100%, 70%) and distilled water. Sections were then incubated for 5 min at room temperature in 0.2% (w/v) aqueous phosphomolybdic acid to suppress non-specific background staining, followed by a brief rinse in distilled water. Sections were subsequently stained for 60 minutes at room temperature with 0.1% Picrosirius Red solution, prepared by dissolving 0.5 g Sirius Red F3B in 500 mL saturated aqueous picric acid. Following staining, sections were washed twice in 0.5% glacial acetic acid in distilled water to remove unbound dye, then dehydrated through an ascending graded ethanol series, and cleared twice in 100% xylene for 2 minutes each. Sections were mounted using Entellan 1079600500, Merck).

With this method, collagen fibers are selectively stained, appearing red under bright-field illumination and exhibiting strong birefringence under polarized light, allowing discrimination of fibrillar collagen organization. Images were acquired using a DM6B microscope (Leica Microsystems) under both bright-field and polarized light conditions.

### Whole-Mount Alkaline Phosphatase Staining

For whole-mount alkaline phosphatase staining, hearts were fixed for 1 hour at room temperature under gentle agitation in 2% formalin/PBS, then washed three times for 10 minutes each in PBS. Samples were subsequently equilibrated in 2 mL alkaline buffer (100 mM Tris-HCl, 100 mM NaCl, 0.1% Tween-20, pH 9.5) for 45 minutes at room temperature.

The alkaline phosphatase reaction was initiated by adding NBT (1.7 µL/mL) and BCIP (1.75 µL/mL) to the alkaline buffer. After 7 minutes of incubation, the chromogenic reaction was stopped by three washes in PBS for 10 minutes each. Samples were imaged immediately upon confirmation of adequate vascular staining. Brightfield images were acquired using an M205 FA stereo microscope (Leica Microsystems).

### Confocal microscopy image acquisition

Confocal images were acquired using a Leica SP5 microscope equipped with diode, argon diode-pumped solid-state, and helium-neon lasers. Depending on the application, one of three objectives was selected: Plan Apo 20×/0.75 multi-immersion, Plan Apo 40×/1.3 oil immersion, or Plan Apo 63×/1.3 glycerol immersion. An electronic zoom factor of 1.5 was applied unless otherwise specified.

Fluorescence excitation was performed at 405, 488, 561, and 633 nm, with emission collected in the following detection windows: 415–480 nm, 498–550 nm, 571–630 nm, and 643–780 nm. Fluorescence signals were detected using hybrid detectors in photon-counting mode or photomultiplier tube (PMT) detectors, as appropriate. For each acquisition, laser power, gain, offset, and detection ranges were individually optimized to prevent inter-channel bleed-through, which was verified prior to imaging. Large-field composite images were assembled by tile scanning and stitching of adjacent fields. BrdU and PCNA colocalization analyses were performed on confocal image stacks acquired as described above.

### Widefield microscopy image acquisition

Histological and fluorescence images were acquired using a fully automated upright Leica DM6B widefield microscope. Fluorescence illumination was provided by a Lumencor SOLA Light Engine, and transmitted light illumination by a Leica CTR6 LED source. Three objectives were available for use depending on the application: Plan Fluotar 10×/0.32 dry, Plan Apochromat 20×/0.8 dry, and Plan Apochromat 40×/0.95 dry.

Fluorescence excitation and emission were achieved using CFP, FITC, Cy3, and Cy5 filter cubes, enabling detection of the corresponding fluorophores. Fluorescence images were captured using either a Hamamatsu ORCA-Fusion sCMOS camera (C14440-20UP) or a Leica DFC9000GT sCMOS camera, selected based on sensitivity requirements. Brightfield images were acquired using a Leica DMC5400 color CMOS camera. The entire system was operated using LAS X Navigator software (Leica Microsystems).

### Stereomicroscope imaging of whole hearts

Whole hearts were imaged using a Leica M205 FA stereomicroscope equipped with a K3 camera at 16× magnification. Samples were positioned in a Petri dish containing 1% solidified agarose to ensure stable mounting, either after fixation in PBS or after cryoprotection in 30% sucrose/PBS. Transmitted light illumination was provided from below, with the angle of incidence manually adjusted to maximize image contrast.

### Image analysis

To ensure representative sampling, multiple tissue sections per heart were imaged and analyzed at each time point. Image processing and quantification were performed using Adobe Photoshop CS6, Fiji/ImageJ, and R. Custom ImageJ macros were developed to automate colocalization and area quantification across multiple regions of interest (ROIs). The key steps of each macro are outlined in the relevant sections below; full scripts are available upon request.

### Definition of regions of interest and ROI area/volume calculation

For each fluorescence channel, background was subtracted using the rolling ball algorithm with a channel-appropriate radius, followed by brightness and contrast adjustment. To optimize particle detection with the "Analyze Particles" function (see below), images were subjected to mild Gaussian blurring to reduce noise, and the watershed algorithm was applied to separate adjoining nuclear signals and to segment the ventricular myocardium, thereby excluding internal lumens and cavities from area measurements.

Using DAPI and muscle staining as references, the entire heart section and wound area were manually delineated. The macro then automatically defined the border zone as a 100 µm-wide band extending from the wound edge into the intact remote myocardium. The areas of all defined ROIs were automatically saved for downstream analysis.

For volume estimation, all sections from each heart were analyzed as described above to obtain the cross-sectional area of the remote myocardium and wound per section. Total volumes were estimated by summing these areas across all sections and multiplying by the effective section interval, calculated as the product of section thickness (12 µm) and the number of slides sampled per heart (8 slides), yielding an interval of 96 µm between analyzed sections. Wound size was expressed as a percentage of total ventricular volume by dividing the wound volume by the total ventricular volume.

### Segmentation and mask extraction

A manual thresholding strategy was applied independently to each fluorescence channel to generate binary masks representing the signal of interest. ROI-specific masks were then extracted from these binary images, and particle analysis was performed on each channel of interest using empirically determined size and circularity parameters.

### Area quantification – L-Plastin, Mpx, Fibronectin

The total area occupied by all detected particles was measured for each channel of interest. For L-Plastin and Mpx, total particle area was normalized to the area of the corresponding ROI to account for differences in section size. For fibronectin, all sections from a single heart were analyzed and an approximate fibronectin volume was calculated as described above, then expressed as a percentage of the total ventricular volume. Each data point represents the mean value across several non-consecutive sections from the same heart, scaled by a factor of 10³ to yield values greater than 1.

### Colocalization analysis – BrdU and PCNA

Colocalization was defined as a minimum 50% areal overlap between a DAPI-segmented nucleus and the binary mask of the channel of interest. Nuclei satisfying this criterion for more than one channel were counted as positive for each. Two metrics were derived: "normalized count in cardiomyocytes (CMs)" represents the number of nuclei positive for all three markers (BrdU⁺/TPM⁺/DAPI⁺ or PCNA⁺/TPM⁺/DAPI⁺) divided by the total number of TPM⁺/DAPI⁺ nuclei, thereby restricting the analysis to the cardiomyocyte population; "normalized count" represents the number of nuclei positive for two markers (BrdU⁺/DAPI⁺ or PCNA⁺/DAPI⁺) divided by the total number of DAPI⁺ nuclei, providing a measure across all cell types. Each data point represents the mean value across several non-consecutive sections from the same heart, scaled by a factor of 10³.

### Quantification of the wound bulging phenotype

Based on AFOG and phalloidin/DAPI staining of heart sections, each ventricle was classified into one of three categories: (1) no or minimal injury, (2) clearly visible injury without bulging, and (3) injury bulging beyond the normal ventricular circumference. To minimize observer bias, two independent analyses were conducted by two different investigators, each performing the full workflow from cryoinjury to image analysis. For one of the two analyses, image classification was performed by a blinded lab member. The results from both analyses were combined for the data presented in this paper.

### RNA extraction, library preparation and sequencing

At 7 days post-cryoinjury (dpci), 24 hearts from uninjured and 24 from cryoinjured fish were collected for each species (zebrafish and platyfish). Ventricles were dissected from the atrium and bulbus arteriosus following a brief incubation with heparin (5 mg/mL) to minimize blood cell contamination. Eight ventricles were pooled per sample, yielding three biological replicates per condition. Samples were snap-frozen on dry ice in Eppendorf tubes containing a single steel bead and stored at −80°C until further processing.

Tissue lysis was performed using a TissueLyser LT (Qiagen) in 75% TRIzol/RNase-free water until complete homogenization was achieved. RNA was isolated and purified using the RNeasy Plus Micro Kit (Qiagen), with on-column DNase digestion (RNase-Free DNase Set, Qiagen) performed to eliminate residual genomic DNA contamination. RNA quantity was assessed using a NanoDrop spectrophotometer and quality was evaluated using an Agilent TapeStation. cDNA was synthesized and amplified using the SMART-Seq Low Input RNA Kit for Sequencing (Takara Bio). RNA-seq libraries were prepared from total RNA using the TruSeq Stranded mRNA Library Preparation Kit (Illumina) and sequenced on an Illumina HiSeq 3000 platform. Raw sequencing data have been deposited in the NCBI Gene Expression Omnibus (GEO) under accession number: <u>GSE305467</u>.

### Raw data processing and differential gene expression analysis

Raw sequencing data quality was assessed using FastQC and RSeQC. Reads were mapped to species-specific reference genomes (*Danio_rerio*.GRCz11.94 for zebrafish and *Xiphophorus_maculatus*.-5.0-male.95 for platyfish) using HiSat2. Read counts per gene were quantified using featureCounts according to the respective genome annotations. All downstream analyses were performed in R.

Differential gene expression (DGE) between experimental groups was tested using the DESeq2 package (Bioconductor). DESeq2-normalized expression values were visualized using volcano plots and scatter plots. Gene ontology (GO) enrichment analysis and gene set enrichment analysis (GSEA) were performed using clusterProfiler, with gene sets derived from KEGG ^113^ and MSigDb ^114^ (**suppl. Table S3**). An interactive Shiny application was developed to facilitate exploration and visualization of the RNA-seq results and is available upon request. Software versions and package references are provided in the Supplementary Data.

### Cross-species RNA-seq analysis

Orthologous gene correspondence between platyfish and zebrafish was established by integrating data from the Ensembl and Orthogene databases, yielding a final dataset of 15,609 one-to-one orthologs (**suppl. Table S1**).

To identify transcripts associated with specific cell types or cellular processes, marker genes were curated from the literature and filtered based on their expression abundance patterns in zebrafish.

### Comparison of transcriptome upon cryoinjury between species

For scatter plot analysis, DESeq2-derived log₂ fold change values were compared across one-to-one orthologous genes between species. A log₂ fold change difference of 1 between species was used as a threshold to highlight genes differentially enriched in response to cryoinjury.

To enable direct cross-species and cross-condition comparison of transcript abundance, DESeq2-normalized counts for each gene were divided by the total normalized counts of the corresponding replicate and multiplied by 10⁶, yielding a TPM-like metric (Transcripts Per Million). This library-size normalization ensures all samples are scaled to a common depth, allowing direct comparison of orthologous gene expression levels across conditions and species. This dataset was merged with the ortholog clustering dataset described above (**suppl. Table S2**). Genes highlighted in the figures were selected based on their normalized expression levels, inter-replicate variability, cross-species and cross-condition differences, and existing knowledge of zebrafish ventricular regeneration.

### Plot and statistical analysis

All plots and statistical analyses were performed using custom R and ImageJ scripts, available upon request. Software versions and package details are provided in the Supplementary Data.

## Supporting information

Sup. Table S1 - clean_combined_wide_orthologue_df

Sup. Table S2 - NRC_clustered_ortho_gene

Sup. Table S3.1 - ZF_gsea_results

Sup. Table S3.1 - PF_gsea_results

## Acknowledgments

We thank V. Zimmermann and Dr. S. Bertho for excellent technical assistance and fish husbandry; Dr. P. Nicholson (University of Bern) for support with RNA sequencing; the Bioimage Core Facility (University of Fribourg) for confocal microscopy support; Dr. C. Pfefferli for assistance with RNA sequencing experiments; and Dr. R. Leech for valuable discussions and critical reading of the manuscript. This study was supported by the Swiss National Science Foundation (SNSF).

## Ethics approval

This study complied with all relevant ethical regulations. Zebrafish were bred, raised, and maintained in accordance with the FELASA guidelines ^115^. The animal housing and all experimental procedures were approved by the cantonal veterinary office of Fribourg, Switzerland.

## Data availability

The authors declare that all data supporting the findings of this study are available within the article and its Supplemental Material files, or from the corresponding author upon request.

## Author’s contributions

Investigations, all authors

Data curation, VH, LR, SB, HL, RB

Formal analyses, VH, LR, HL, RB

Conceptualization, AJ, SB

Visualization, VH, LR, SB

Supervision, AJ, SB, RB

Writing - original draft, VH, AJ

Writing - review & editing, all authors

## Declaration of generative AI and AI-assisted technologies in the manuscript preparation process

During the preparation of this work, the author(s) used Perplexity AI and Claude AI to correct the English grammar and syntax. After using this tool/service, the author(s) reviewed and edited the content as needed and take(s) full responsibility for the content of the published article.

## References

1. Saleh, M., and Ambrose, J.A. (2018). Understanding myocardial infarction. F1000Res *7*. 10.12688/f1000research.15096.1.

2. Prabhu, S.D., and Frangogiannis, N.G. (2016). The Biological Basis for Cardiac Repair After Myocardial Infarction. Circ Res 119, 91–112. 10.1161/circresaha.116.303577.

3. Poss, K.D., Wilson, L.G., and Keating, M.T. (2002). Heart regeneration in zebrafish. Science 298, 2188–2190.

4. Flink, I.L. (2002). Cell cycle reentry of ventricular and atrial cardiomyocytes and cells within the epicardium following amputation of the ventricular apex in the axolotl, Amblystoma mexicanum: confocal microscopic immunofluorescent image analysis of bromodeoxyuridine-labeled nuclei. Anat Embryol (Berl) 205, 235–244. 10.1007/s00429-002-0249-6.

5. Jopling, C., Sleep, E., Raya, M., Marti, M., Raya, A., and Izpisua Belmonte, J.C. (2010). Zebrafish heart regeneration occurs by cardiomyocyte dedifferentiation and proliferation. Nature 464, 606–609. 10.1038/nature08899.

6. Kikuchi, K., Holdway, J.E., Werdich, A.A., Anderson, R.M., Fang, Y., Egnaczyk, G.F., Evans, T., Macrae, C.A., Stainier, D.Y., and Poss, K.D. (2010). Primary contribution to zebrafish heart regeneration by gata4(+) cardiomyocytes. Nature 464, 601–605. 10.1038/nature08804.

7. Cano-Martínez, A., Vargas-González, A., Guarner-Lans, V., Prado-Zayago, E., León-Oleda, M., and Nieto-Lima, B. (2010). Functional and structural regeneration in the axolotl heart (Ambystoma mexicanum) after partial ventricular amputation. Arch Cardiol Mex 80, 79–86.

8. Witman, N., Murtuza, B., Davis, B., Arner, A., and Morrison, J.I. (2011). Recapitulation of developmental cardiogenesis governs the morphological and functional regeneration of adult newt hearts following injury. Dev Biol 354, 67–76. 10.1016/j.ydbio.2011.03.021.

9. Pronobis, M.I., and Poss, K.D. (2020). Signals for cardiomyocyte proliferation during zebrafish heart regeneration. Curr Opin Physiol 14, 78–85. 10.1016/j.cophys.2020.02.002.

10. Huang, H., Huang, G.N., and Payumo, A.Y. (2024). Two decades of heart regeneration research: Cardiomyocyte proliferation and beyond. WIREs Mech Dis 16, e1629. 10.1002/wsbm.1629.

11. Weinberger, M., and Riley, P.R. (2024). Animal models to study cardiac regeneration. Nature Reviews Cardiology 21, 89–105. 10.1038/s41569-023-00914-x.

12. Mehdipour, M., Park, S., and Huang, G.N. (2023). Unlocking cardiomyocyte renewal potential for myocardial regeneration therapy. J Mol Cell Cardiol 177, 9–20. 10.1016/j.yjmcc.2023.02.002.

13. Jensen, B., Wang, T., Christoffels, V.M., and Moorman, A.F. (2013). Evolution and development of the building plan of the vertebrate heart. Biochim Biophys Acta 1833, 783–794. 10.1016/j.bbamcr.2012.10.004.

14. Betancur-R, R., Wiley, E.O., Arratia, G., Acero, A., Bailly, N., Miya, M., Lecointre, G., and Ortí, G. (2017). Phylogenetic classification of bony fishes. BMC evolutionary biology 17, 162–162. 10.1186/s12862-017-0958-3.

15. Hughes, L.C., Ortí, G., Huang, Y., Sun, Y., Baldwin, C.C., Thompson, A.W., Arcila, D., Betancur, R.R., Li, C., Becker, L., et al. (2018). Comprehensive phylogeny of ray-finned fishes (Actinopterygii) based on transcriptomic and genomic data. Proc Natl Acad Sci U S A 115, 6249–6254. 10.1073/pnas.1719358115.

16. Clarke, J.T., Lloyd, G.T., and Friedman, M. (2016). Little evidence for enhanced phenotypic evolution in early teleosts relative to their living fossil sister group. Proc Natl Acad Sci U S A 113, 11531–11536. 10.1073/pnas.1607237113.

17. Santini, F., Harmon, L.J., Carnevale, G., and Alfaro, M.E. (2009). Did genome duplication drive the origin of teleosts? A comparative study of diversification in ray-finned fishes. BMC Evolutionary Biology 9, 194. 10.1186/1471-2148-9-194.

18. McWilliam, J.A. (1885). On the Structure and Rhythm of the Heart in Fishes, with especial reference to the Heart of the Eel. The Journal of Physiology 6, 192–292. 10.1113/jphysiol.1885.sp000196.

19. Santer, R.M. (1985). Morphology and innervation of the fish heart. (Springer-Verlag).

20. Chen, X., Zhong, X., and Huang, G.N. (2024). Heart regeneration from the whole-organism perspective to single-cell resolution. npj Regenerative Medicine 9, 34. 10.1038/s41536-024-00378-8.

21. Dittrich, A., Hansen, K., Simonsen, M.I.T., Busk, M., Alstrup, A.K.O., and Lauridsen, H. (2020). Intrinsic Heart Regeneration in Adult Vertebrates May be Strictly Limited to Low-Metabolic Ectotherms. BioEssays 42, 2000054. 10.1002/bies.202000054.

22. Jaźwińska, A., and Blanchoud, S. (2020). Towards deciphering variations of heart regeneration in fish. Current Opinion in Physiology 14, 21–26. 10.1016/j.cophys.2019.11.007.

23. Meijler, F.L., and Meijler, T.D. (2011). Archetype, adaptation and the mammalian heart. Neth Heart J 19, 142–148 10.1007/s12471-011-0086-4.

24. González-Rosa, J.M., Burns, C.E., and Burns, C.G. (2017). Zebrafish heart regeneration: 15 years of discoveries. Regeneration 4, 105–123. 10.1002/reg2.83.

25. Ross Stewart, K.M., Walker, S.L., Baker, A.H., Riley, P.R., and Brittan, M. (2022). Hooked on heart regeneration: the zebrafish guide to recovery. Cardiovasc Res 118, 1667–1679. 10.1093/cvr/cvab214.

26. Zuppo, D.A., and Tsang, M. (2020). Zebrafish heart regeneration: Factors that stimulate cardiomyocyte proliferation. Semin Cell Dev Biol 100, 3–10. 10.1016/j.semcdb.2019.09.005.

27. Chablais, F., Veit, J., Rainer, G., and Jaźwińska, A. (2011). The zebrafish heart regenerates after cryoinjury-induced myocardial infarction. BMC Developmental Biology 11, 21. 10.1186/1471-213X-11-21.

28. Schnabel, K., Wu, C.C., Kurth, T., and Weidinger, G. (2011). Regeneration of cryoinjury induced necrotic heart lesions in zebrafish is associated with epicardial activation and cardiomyocyte proliferation. PLoS One 6, e18503. 10.1371/journal.pone.0018503.

29. Gonzalez-Rosa, J.M., Martin, V., Peralta, M., Torres, M., and Mercader, N. (2011). Extensive scar formation and regression during heart regeneration after cryoinjury in zebrafish. Development 138, 1663–1674. 10.1242/dev.060897.

30. Bise, T., Sallin, P., Pfefferli, C., and Jaźwińska, A. (2020). Multiple cryoinjuries modulate the efficiency of zebrafish heart regeneration. Scientific Reports 10, 11551. 10.1038/s41598-020-68200-1.

31. Fernandez, C.E., Bakovic, M., and Karra, R. (2018). Endothelial Contributions to Zebrafish Heart Regeneration. J Cardiovasc Dev Dis 5 10.3390/jcdd5040056.

32. Cao, J., and Poss, K.D. (2018). The epicardium as a hub for heart regeneration. Nature Reviews Cardiology 15, 631–647. 10.1038/s41569-018-0046-4.

33. Feng, X., Travisano, S., Pearson, C.A., Lien, C.L., and Harrison, M.R.M. (2021). The Lymphatic System in Zebrafish Heart Development, Regeneration and Disease Modeling. J Cardiovasc Dev Dis 8 10.3390/jcdd8020021.

34. Ryan, R., Moyse, B.R., and Richardson, R.J. (2020). Zebrafish cardiac regeneration—looking beyond cardiomyocytes to a complex microenvironment. Histochemistry and Cell Biology 154, 533–548. 10.1007/s00418-020-01913-6.

35. Simões, F.C., and Riley, P.R. (2022). Immune cells in cardiac repair and regeneration. Development 149. 10.1242/dev.199906.

36. Mahmoud, Ahmed I., O’Meara, Caitlin C., Gemberling, M., Zhao, L., Bryant, Donald M., Zheng, R., Gannon, Joseph B., Cai, L., Choi, W.-Y., Egnaczyk, Gregory F., et al. (2015). Nerves Regulate Cardiomyocyte Proliferation and Heart Regeneration. Developmental Cell 34, 387–399. 10.1016/j.devcel.2015.06.017.

37. Ito, K., Morioka, M., Kimura, S., Tasaki, M., Inohaya, K., and Kudo, A. (2014). Differential reparative phenotypes between zebrafish and medaka after cardiac injury. Developmental Dynamics 243, 1106–1115. 10.1002/dvdy.24154.

38. Potts, H.G., Stockdale, W.T., and Mommersteeg, M.T.M. (2021). Unlocking the Secrets of the Regenerating Fish Heart: Comparing Regenerative Models to Shed Light on Successful Regeneration. Journal of Cardiovascular Development and Disease 8, 4.

39. Lai, S.-L., Marín-Juez, R., Moura, P.L., Kuenne, C., Lai, J.K.H., Tsedeke, A.T., Guenther, S., Looso, M., and Stainier, D.Y.R. (2017). Reciprocal analyses in zebrafish and medaka reveal that harnessing the immune response promotes cardiac regeneration. eLife 6. 10.7554/eLife.25605.

40. Tiozzo, S., and Copley, R.R. (2015). Reconsidering regeneration in metazoans: an evo-devo approach. Frontiers in Ecology and Evolution Volume 3 - 2015. 10.3389/fevo.2015.00067.

41. Fumagalli, M.R., Zapperi, S., and La Porta, C.A. (2018). Regeneration in distantly related species: common strategies and pathways. NPJ Systems Biology and Applications 4, 1–8.

42. Elchaninov, A., Sukhikh, G., and Fatkhudinov, T. (2021). Evolution of Regeneration in Animals: A Tangled Story. Frontiers in Ecology and Evolution 9. 10.3389/fevo.2021.621686.

43. Dornburg, A., and Near, T.J. (2021). The Emerging Phylogenetic Perspective on the Evolution of Actinopterygian Fishes. Annual Review of Ecology, Evolution, and Systematics 52, 427–452. 10.1146/annurev-ecolsys-122120-122554.

44. Parey, E., Louis, A., Montfort, J., Bouchez, O., Roques, C., Iampietro, C., Lluch, J., Castinel, A., Donnadieu, C., Desvignes, T., et al. (2023). Genome structures resolve the early diversification of teleost fishes. Science 379, 572–575. doi:10.1126/science.abq4257.

45. Schartl, M., and Lu, Y. (2024). Validity of Xiphophorus fish as models for human disease. Dis Model Mech 17. 10.1242/dmm.050382.

46. Schartl, M., Kneitz, S., Ormanns, J., Schmidt, C., Anderson, J.L., Amores, A., Catchen, J., Wilson, C., Geiger, D., Du, K., et al. (2021). The Developmental and Genetic Architecture of the Sexually Selected Male Ornament of Swordtails. Current Biology 31, 911–922.e914. 10.1016/j.cub.2020.11.028.

47. Tobler, M., Kelley, J.L., Plath, M., and Riesch, R. (2018). Extreme environments and the origins of biodiversity: Adaptation and speciation in sulphide spring fishes. Molecular Ecology 27, 843–859. 10.1111/mec.14497.

48. Schartl, M., and Walter, R.B. (2016). Xiphophorus and Medaka Cancer Models. In Cancer and Zebrafish: Mechanisms, Techniques, and Models, D.M. Langenau, ed. (Springer International Publishing), pp. 531–552. 10.1007/978-3-319-30654-4_23.

49. Safian, D., Wiegertjes, G.F., and Pollux, B.J.A. (2021). The Fish Family Poeciliidae as a Model to Study the Evolution and Diversification of Regenerative Capacity in Vertebrates. Frontiers in Ecology and Evolution Volume 9 - 2021. 10.3389/fevo.2021.613157.

50. Pollux, B.J.A., Pires, M.N., Banet, A.I., and Reznick, D.N. (2009). Evolution of Placentas in the Fish Family Poeciliidae: An Empirical Study of Macroevolution. Annual Review of Ecology, Evolution, and Systematics 40, 271–289. 10.1146/annurev.ecolsys.110308.120209.

51. Schartl, M., Walter, R.B., Shen, Y., Garcia, T., Catchen, J., Amores, A., Braasch, I., Chalopin, D., Volff, J.N., Lesch, K.P., et al. (2013). The genome of the platyfish, Xiphophorus maculatus, provides insights into evolutionary adaptation and several complex traits. Nat Genet 45, 567–572. 10.1038/ng.2604.

52. Rees, L., König, D., and Jaźwińska, A. (2022). Platyfish bypass the constraint of the caudal fin ventral identity in teleosts. Developmental Dynamics 251, 1862–1879. 10.1002/dvdy.518.

53. Rees, L., König, D., and Jaźwińska, A. (2023). Regeneration of the dermal skeleton and wound epidermis formation depend on BMP signaling in the caudal fin of platyfish. Froniers in Cell and Developmental Biology 11. 10.3389/fcell.2023.1134451.

54. Sallin, P., de Preux Charles, A.S., Duruz, V., Pfefferli, C., and Jaźwińska, A. (2015). A dual epimorphic and compensatory mode of heart regeneration in zebrafish. Dev Biol 399, 27–40. 10.1016/j.ydbio.2014.12.002.

55. Pfefferli, C., and Jaźwińska, A. (2017). The careg element reveals a common regulation of regeneration in the zebrafish myocardium and fin. Nature Communications 8, 15151. 10.1038/ncomms15151.

56. Hu, N., Yost, H.J., and Clark, E.B. (2001). Cardiac morphology and blood pressure in the adult zebrafish. Anat Rec 264, 1–12.

57. Icardo, J.M. (2013). Collagen and elastin histochemistry of the teleost bulbus arteriosus: False positives. Acta Histochemica 115, 185–189. 10.1016/j.acthis.2012.03.002.

58. Kammerer, J., Cirnu, A., Williams, T., Hasselmeier, M., Nörpel, M., Chen, R., and Gerull, B. (2024). Macro-based collagen quantification and segmentation in picrosirius red-stained heart sections using light microscopy. Biology Methods and Protocols 9, bpae027. 10.1093/biomethods/bpae027.

59. Herwig, L., Blum, Y., Krudewig, A., Ellertsdottir, E., Lenard, A., Belting, H.G., and Affolter, M. (2011). Distinct cellular mechanisms of blood vessel fusion in the zebrafish embryo. Curr Biol 21, 1942–1948. 10.1016/j.cub.2011.10.016.

60. Chablais, F., and Jaźwińska, A. (2012). Induction of myocardial infarction in adult zebrafish using cryoinjury. Journal of Visualized Experiments (JoVE), e3666. 10.3791/3666.

61. González-Rosa, J.M., Martín, V., Peralta, M., Torres, M., and Mercader, N. (2011). Extensive scar formation and regression during heart regeneration after cryoinjury in zebrafish. Development 138, 1663–1674. 10.1242/dev.060897.

62. Pfefferli, C., Bonvin, M., Grepper, D., Robatel, S., König, D., Lischer, H.E.L., Bruggmann, R., and Jaźwińska, A. (2023). Parallels between oncogene-driven cardiac hyperplasia and heart regeneration in zebrafish. Development 150. 10.1242/dev.201412.

63. González-Rosa, J.M., Sharpe, M., Field, D., Soonpaa, M.H., Field, L.J., Burns, C.E., and Burns, C.G. (2018). Myocardial Polyploidization Creates a Barrier to Heart Regeneration in Zebrafish. Developmental Cell 44, 433–446.e437. 10.1016/j.devcel.2018.01.021.

64. Ryan, R., Moyse, B.R., and Richardson, R.J. (2020). Zebrafish cardiac regeneration-looking beyond cardiomyocytes to a complex microenvironment. Histochem Cell Biol 154, 533–548. 10.1007/s00418-020-01913-6.

65. Carey, C.M., Hollins, H.L., Schmid, A.V., and Gagnon, J.A. (2023). Distinct features of the regenerating heart uncovered through comparative single-cell profiling. bioRxiv. 10.1101/2023.07.04.547574.

66. Missinato, M.A., Saydmohammed, M., Zuppo, D.A., Rao, K.S., Opie, G.W., Kühn, B., and Tsang, M. (2018). Dusp6 attenuates Ras/MAPK signaling to limit zebrafish heart regeneration. Development 145. 10.1242/dev.157206.

67. Zhao, L., Ben-Yair, R., Burns, C.E., and Burns, C.G. (2019). Endocardial Notch Signaling Promotes Cardiomyocyte Proliferation in the Regenerating Zebrafish Heart through Wnt Pathway Antagonism. Cell Reports 26, 546–554.e545. 10.1016/j.celrep.2018.12.048.

68. Wang, W., Hu, Y.F., Pang, M., Chang, N., Yu, C., Li, Q., Xiong, J.W., Peng, Y., and Zhang, R. (2021). BMP and Notch Signaling Pathways differentially regulate Cardiomyocyte Proliferation during Ventricle Regeneration. Int J Biol Sci 17, 2157–2166. 10.7150/ijbs.59648.

69. Chablais, F., and Jaźwińska, A. (2012). The regenerative capacity of the zebrafish heart is dependent on TGFbeta signaling. Development 139, 1921–1930. 10.1242/dev.078543.

70. Kanwal, Z., Wiegertjes, G.F., Veneman, W.J., Meijer, A.H., and Spaink, H.P. (2014). Comparative studies of Toll-like receptor signalling using zebrafish. Dev Comp Immunol 46, 35–52. 10.1016/j.dci.2014.02.003.

71. Bongiovanni, C., Bueno-Levy, H., Posadas Pena, D., Del Bono, I., Miano, C., Boriati, S., Da Pra, S., Sacchi, F., Redaelli, S., Bergen, M., et al. (2024). BMP7 promotes cardiomyocyte regeneration in zebrafish and adult mice. Cell Rep 43, 114162. 10.1016/j.celrep.2024.114162.

72. Rasouli, S.J., and Stainier, D.Y.R. (2017). Regulation of cardiomyocyte behavior in zebrafish trabeculation by Neuregulin 2a signaling. Nat Commun 8, 15281. 10.1038/ncomms15281.

73. de Bakker, D.E.M., Bouwman, M., Dronkers, E., Simões, F.C., Riley, P.R., Goumans, M.J., Smits, A.M., and Bakkers, J. (2021). Prrx1b restricts fibrosis and promotes Nrg1-dependent cardiomyocyte proliferation during zebrafish heart regeneration. Development 148. 10.1242/dev.198937.

74. Redd, M.J., Kelly, G., Dunn, G., Way, M., and Martin, P. (2006). Imaging macrophage chemotaxis in vivo: Studies of microtubule function in zebrafish wound inflammation. Cell Motility and the Cytoskeleton 63, 415–422. 10.1002/cm.20133.

75. Morley, S.C. (2012). The actin-bundling protein L-plastin: a critical regulator of immune cell function. Int J Cell Biol 2012, 935173. 10.1155/2012/935173.

76. Keightley, M.-C., Wang, C.-H., Pazhakh, V., and Lieschke, G.J. (2014). Delineating the roles of neutrophils and macrophages in zebrafish regeneration models. The International Journal of Biochemistry & Cell Biology 56, 92–106. 10.1016/j.biocel.2014.07.010.

77. Simões, F.C., Cahill, T.J., Kenyon, A., Gavriouchkina, D., Vieira, J.M., Sun, X., Pezzolla, D., Ravaud, C., Masmanian, E., Weinberger, M., et al. (2020). Macrophages directly contribute collagen to scar formation during zebrafish heart regeneration and mouse heart repair. Nature Communications 11, 600. 10.1038/s41467-019-14263-2.

78. Bevan, L., Lim, Z.W., Venkatesh, B., Riley, P.R., Martin, P., and Richardson, R.J. (2019). Specific macrophage populations promote both cardiac scar deposition and subsequent resolution in adult zebrafish. Cardiovascular Research 116, 1357–1371. 10.1093/cvr/cvz221.

79. Wei, K.H., Lin, I.T., Chowdhury, K., Lim, K.L., Liu, K.T., Ko, T.M., Chang, Y.M., Yang, K.C., and Lai, S.B. (2023). Comparative single-cell profiling reveals distinct cardiac resident macrophages essential for zebrafish heart regeneration. Elife 12. 10.7554/eLife.84679.

80. Beisaw, A., Kuenne, C., Guenther, S., Dallmann, J., Wu, C.-C., Bentsen, M., Looso, M., and Stainier, D.Y.R. (2020). AP-1 Contributes to Chromatin Accessibility to Promote Sarcomere Disassembly and Cardiomyocyte Protrusion During Zebrafish Heart Regeneration. Circ Res 126, 1760–1778. 10.1161/CIRCRESAHA.119.316167.

81. Tamaki, T., Yoshida, T., Shibata, E., Nishihara, H., Ochi, H., and Kawakami, A. (2023). Splashed E-box and AP-1 motifs cooperatively drive regeneration response and shape regeneration abilities. Biology Open 12, bio059810. 10.1242/bio.059810.

82. Xiao, C., Gao, L., Hou, Y., Xu, C., Chang, N., Wang, F., Hu, K., He, A., Luo, Y., Wang, J., et al. (2016). Chromatin-remodelling factor Brg1 regulates myocardial proliferation and regeneration in zebrafish. Nature Communications 7, 13787. 10.1038/ncomms13787.

83. Hughes, S.M., Cho, M., Karsch-Mizrachi, I., Travis, M., Silberstein, L., Leinwand, L.A., and Blau, H.M. (1993). Three Slow Myosin Heavy Chains Sequentially Expressed in Developing Mammalian Skeletal Muscle. Developmental Biology 158, 183–199. 10.1006/dbio.1993.1178.

84. Maggs, A.M., Taylor-Harris, P., Peckham, M., and Hughes, S.M. (2000). Evidence for differential post-translational modifications of slow myosin heavy chain during murine skeletal muscle development. Journal of Muscle Research & Cell Motility 21, 101–113. 10.1023/A:1005639229497.

85. Yin, V.P., Lepilina, A., Smith, A., and Poss, K.D. (2012). Regulation of zebrafish heart regeneration by miR-133. Developmental Biology 365, 319–327. 10.1016/j.ydbio.2012.02.018.

86. Sánchez-Iranzo, H., Galardi-Castilla, M., Minguillón, C., Sanz-Morejón, A., González-Rosa, J.M., Felker, A., Ernst, A., Guzmán-Martínez, G., Mosimann, C., and Mercader, N. (2018). Tbx5a lineage tracing shows cardiomyocyte plasticity during zebrafish heart regeneration. Nature Communications 9. 10.1038/s41467-017-02650-6.

87. Forman-Rubinsky, R., Paul, A., Feng, W., Schlegel, B.T., Zuppo, D.A., Kedziora, K., Stoltz, D.B., Watkins, S.C., Rajasundaram, D., Li, G., and Tsang, M. (2025). Cited4a limits cardiomyocyte dedifferentiation and proliferation during zebrafish heart regeneration. Development 152. 10.1242/dev.204659.

88. Huang, M., Akerberg, A.A., Zhang, X., Yoon, H., Joshi, S., Hallinan, C., Nguyen, C., Pu, W.T., Haigis, M.C., Burns, C.G., and Burns, C.E. (2022). Intrinsic myocardial defects underlie an Rbfox-deficient zebrafish model of hypoplastic left heart syndrome. Nature Communications 13, 5877. 10.1038/s41467-022-32982-x.

89. Huang, M., Ladha, F.A., Wang, Y., Trembley, M.A., Yin, H.-M., Zheng, R., Prondzynski, M., Tharani, Y., Akerberg, A.A., Aigner, S., et al. (2025). Dosage-sensitive *RBFOX2* autoregulation promotes cardiomyocyte differentiation by maturing the transcriptome. bioRxiv, 2025.2010.2028.685214. 10.1101/2025.10.28.685214.

90. Frese, K.S., Meder, B., Keller, A., Just, S., Haas, J., Vogel, B., Fischer, S., Backes, C., Matzas, M., Köhler, D., et al. (2015). RNA splicing regulated by RBFOX1 is essential for cardiac function in zebrafish. Journal of Cell Science 128, 3030–3040. 10.1242/jcs.166850.

91. Vasudevarao, M.D., Posadas Pena, D., Ihle, M., Bongiovanni, C., Maity, P., Geissler, D., Mohammadi, H.F., Rall-Scharpf, M., Niemann, J., Mommersteeg, M.T.M., et al. (2025). BMP signaling promotes zebrafish heart regeneration via alleviation of replication stress. Nature Communications 16, 1708. 10.1038/s41467-025-56993-6.

92. Braasch, I., Peterson, S.M., Desvignes, T., McCluskey, B.M., Batzel, P., and Postlethwait, J.H. (2015). A new model army: Emerging fish models to study the genomics of vertebrate Evo-Devo. Journal of Experimental Zoology Part B: Molecular and Developmental Evolution 324, 316–341. 10.1002/jez.b.22589.

93. Chowdhury, K., Lin, S., and Lai, S.-L. (2022). Comparative Study in Zebrafish and Medaka Unravels the Mechanisms of Tissue Regeneration. Frontiers in Ecology and Evolution Volume 10 - 2022. 10.3389/fevo.2022.783818.

94. Sánchez-Iranzo, H., Galardi-Castilla, M., Sanz-Morejón, A., González-Rosa, J.M., Costa, R., Ernst, A., Sainz de Aja, J., Langa, X., and Mercader, N. (2018). Transient fibrosis resolves via fibroblast inactivation in the regenerating zebrafish heart. Proceedings of the National Academy of Sciences 115, 4188–4193. 10.1073/pnas.1716713115.

95. Marro, J., Pfefferli, C., de Preux Charles, A.S., Bise, T., and Jaźwińska, A. (2016). Collagen XII Contributes to Epicardial and Connective Tissues in the Zebrafish Heart during Ontogenesis and Regeneration. PLoS One 11, e0165497. 10.1371/journal.pone.0165497.

96. Lafontant, P.J., Behzad, A.R., Brown, E., Landry, P., Hu, N., and Burns, A.R. (2013). Cardiac myocyte diversity and a fibroblast network in the junctional region of the zebrafish heart revealed by transmission and serial block-face scanning electron microscopy. PLoS One 8, e72388. 10.1371/journal.pone.0072388.

97. Klaourakis, K., Vieira, J.M., and Riley, P.R. (2021). The evolving cardiac lymphatic vasculature in development, repair and regeneration. Nat Rev Cardiol 18, 368–379. 10.1038/s41569-020-00489-x.

98. Pfefferli, C., and Jaźwińska, A. (2019). Lymphatic vessels help mend broken hearts. eLife 8, e52200. 10.7554/eLife.52200.

99. Kapuria, S., Yoshida, T., and Lien, C.L. (2018). Coronary Vasculature in Cardiac Development and Regeneration. J Cardiovasc Dev Dis 5. 10.3390/jcdd5040059.

100. Lowe, V., Wisniewski, L., and Pellet-Many, C. (2021). The Zebrafish Cardiac Endothelial Cell-Roles in Development and Regeneration. J Cardiovasc Dev Dis 8. 10.3390/jcdd8050049.

101. Stockdale, W.T., Lemieux, M.E., Killen, A.C., Zhao, J., Hu, Z., Riepsaame, J., Hamilton, N., Kudoh, T., Riley, P.R., van Aerle, R., et al. (2018). Heart Regeneration in the Mexican Cavefish. Cell Reports 25, 1997–2007.e1997. 10.1016/j.celrep.2018.10.072.

102. Offen, N., Blum, N., Meyer, A., and Begemann, G. (2008). Fgfr1 signalling in the development of a sexually selected trait in vertebrates, the sword of swordtail fish. BMC Developmental Biology 8. 10.1186/1471-213x-8-98.

103. Patel, S., Ranadive, I., Desai, I., and Balakrishnan, S. (2019). Regeneration of caudal fin in Poecilia latipinna: Insights into the progressive tissue morphogenesis. Organogenesis 15, 35–42.

104. Darnet, S., Dragalzew, A.C., Amaral, D.B., Sousa, J.F., Thompson, A.W., Cass, A.N., Lorena, J., Pires, E.S., Costa, C.M., Sousa, M.P., et al. (2019). Deep evolutionary origin of limb and fin regeneration. Proceedings of the National Academy of Sciences 116, 15106. 10.1073/pnas.1900475116.

105. Cobham, A.E., Kenzior, A., Morales-Sosa, P., Javier, J.E., Swanson, S., Wood, C., and Rohner, N. (2025). Cave adaptation favors aging resilience in the Mexican tetra. npj Metabolic Health and Disease 3, 33. 10.1038/s44324-025-00069-y.

106. Farrell, A.P., Farrell, N.D., Jourdan, H., and Cox, G.K. (2012). A Perspective on the evolution of the coronary circulation in fishes and the transition to terrestrial life. In Ontogeny and Phylogeny of the Vertebrate Heart, S. D., and T. Wang, eds. (Springer), pp. 75–102.

107. Farrell, A.P. (2023). Getting to the heart of anatomical diversity and phenotypic plasticity: fish hearts are an optimal organ model in need of greater mechanistic study. J Exp Biol 226. 10.1242/jeb.245582.

108. Farrell, A.P., and Jones, D.R. (1992). The heart. In Fish Physiology: The Cardiovascular System., W.S. Hoar, D.J. Randall, and A.P. Farrell, eds. (Academic Press), pp. 1–88.

109. Frances, C., Romero, A., and Grady, D. (1998). Left ventricular pseudoaneurysm. Journal of the American College of Cardiology 32, 557–561. 10.1016/s0735-1097(98)00290-3.

110. Soni, M.K., and Raja, S.G. (2020). Post-infarction Ventricular Aneurysms. In Cardiac Surgery, pp. 243–252. 10.1007/978-3-030-24174-2_26.

111. Bise, T., and Jaźwińska, A. (2019). Intrathoracic Injection for the Study of Adult Zebrafish Heart. Journal of Visualized Experiments (JoVE), e59724. doi:10.3791/59724.

112. Oudhoff, H., Hisler, V., Baumgartner, F., Rees, L., Grepper, D., and Jaźwińska, A. (2024). Skeletal muscle regeneration after extensive cryoinjury of caudal myomeres in adult zebrafish. NPJ Regen Med 9, 8. 10.1038/s41536-024-00351-5.

113. Kanehisa, M., Sato, Y., and Kawashima, M. (2022). KEGG mapping tools for uncovering hidden features in biological data. Protein Science 31, 47–53. 10.1002/pro.4172.

114. Liberzon, A., Birger, C., Thorvaldsdóttir, H., Ghandi, M., Mesirov, J.P., and Tamayo, P. (2015). The Molecular Signatures Database (MSigDB) hallmark gene set collection. Cell Syst 1, 417–425. 10.1016/j.cels.2015.12.004.

115. Aleström, P., D’Angelo, L., Midtlyng, P.J., Schorderet, D.F., Schulte-Merker, S., Sohm, F., and Warner, S. (2019). Zebrafish: Housing and husbandry recommendations. Laboratory Animals 54, 213–224. 10.1177/0023677219869037.

116. Nelson, G. (1989). Phylogeny of major fish groups. The hierarchy of life, 325-336.

117. Sanciangco, M.D., Carpenter, K.E., and Betancur, R.R. (2016). Phylogenetic placement of enigmatic percomorph families (Teleostei: Percomorphaceae). Mol Phylogenet Evol 94, 565–576. 10.1016/j.ympev.2015.10.006.

118. Arun Gopinathan, P., Kokila, G., Jyothi, M., Ananjan, C., Pradeep, L., and Humaira Nazir, S. (2015). Study of Collagen Birefringence in Different Grades of Oral Squamous Cell Carcinoma Using Picrosirius Red and Polarized Light Microscopy. Scientifica 2015, 802980. 10.1155/2015/802980.

