## Supplemental figure and material for "The livebearers platyfish and swordtails partially regenerate their hearts with persistent scarring"

**The PDF supplementary data include:**

Supplementary Figures S1 to S8 (pages 2-12)

Supplementary Materials (pages 13-17)

**Separate excel files online:**

Supplementary Table S1 : clean_combined_wide_orthologue_df
Supplementary Table S2 : NRC_clustered_ortho_gene
Supplementary Table S3 : gsea_results

### Supplementary Figure S1

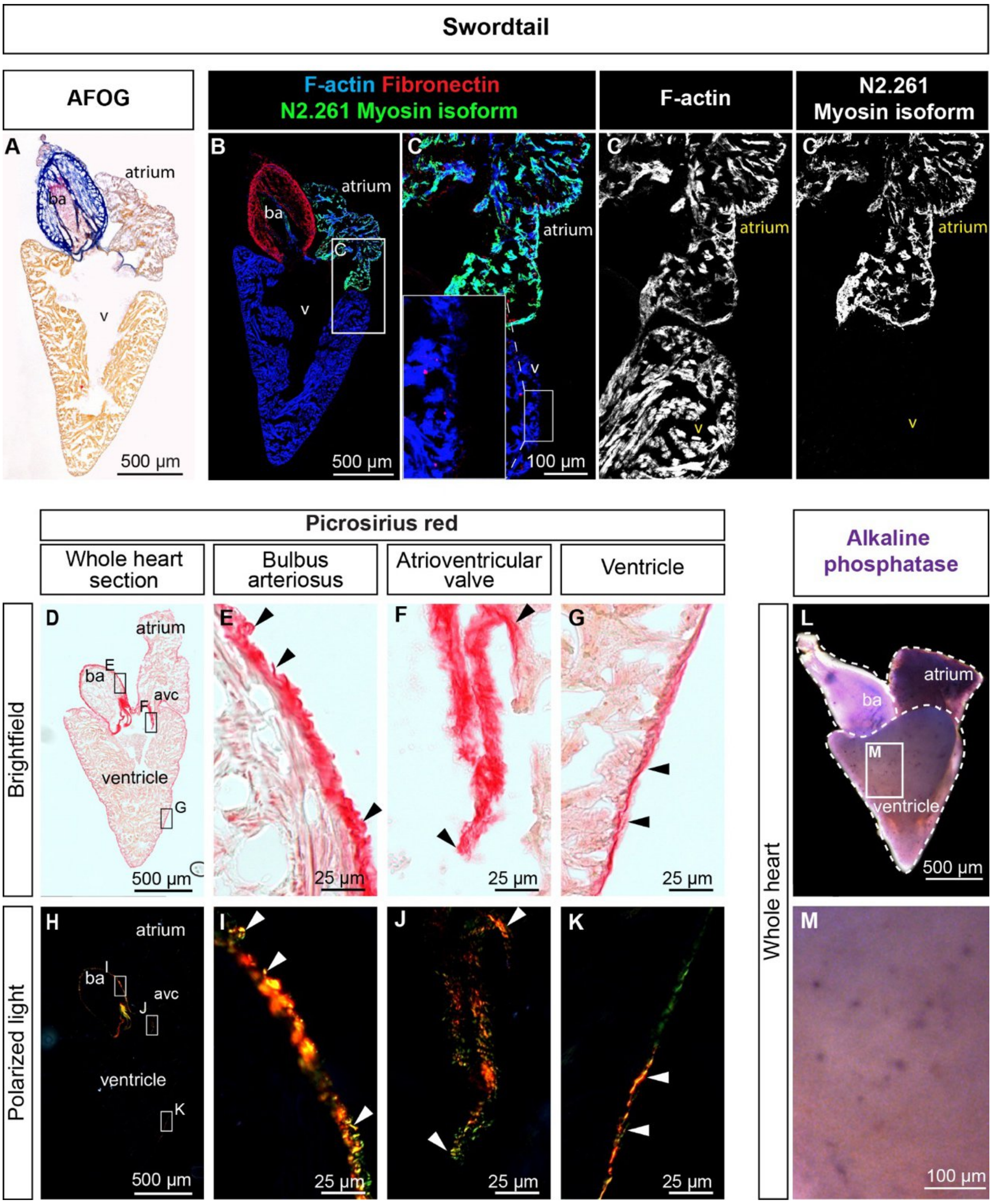


**Suppl. Figure S1. The anatomy of swordtail heart is similar to that of platyfish.**
**(A)** AFOG staining of longitudinal swordtail heart sections. Intact myocardium appears orange; fibrin and protein deposits appear red; collagen appears blue.
**(B–C)** Fluorescence staining of longitudinal swordtail heart sections. F-actin labeling (blue) reveals the absence of a compact myocardial layer, as shown in the magnified inset. The N2.261 antibody (green) immunolabels the atrium, consistent with atrial myosin recognition in swordtails, as observed in platyfish (Figure 1). The bulbus arteriosus is fibronectin-positive (red).
**(D–K)** PSR staining of longitudinal sections imaged under bright-field (D–G) and polarized light (H–K). Frames indicate regions magnified in adjacent panels, labeled with the corresponding letter, showing the bulbus arteriosus, valves, and outer myocardial layer, as in Figure 2. ba, bulbus arteriosus; v, ventricle.
**(L, M)** Alkaline phosphatase staining of the whole heart of the swordtail reveals a complete absence of a coronary vasculature, similar to that observed in platyfish.

### Supplementary Figure S2

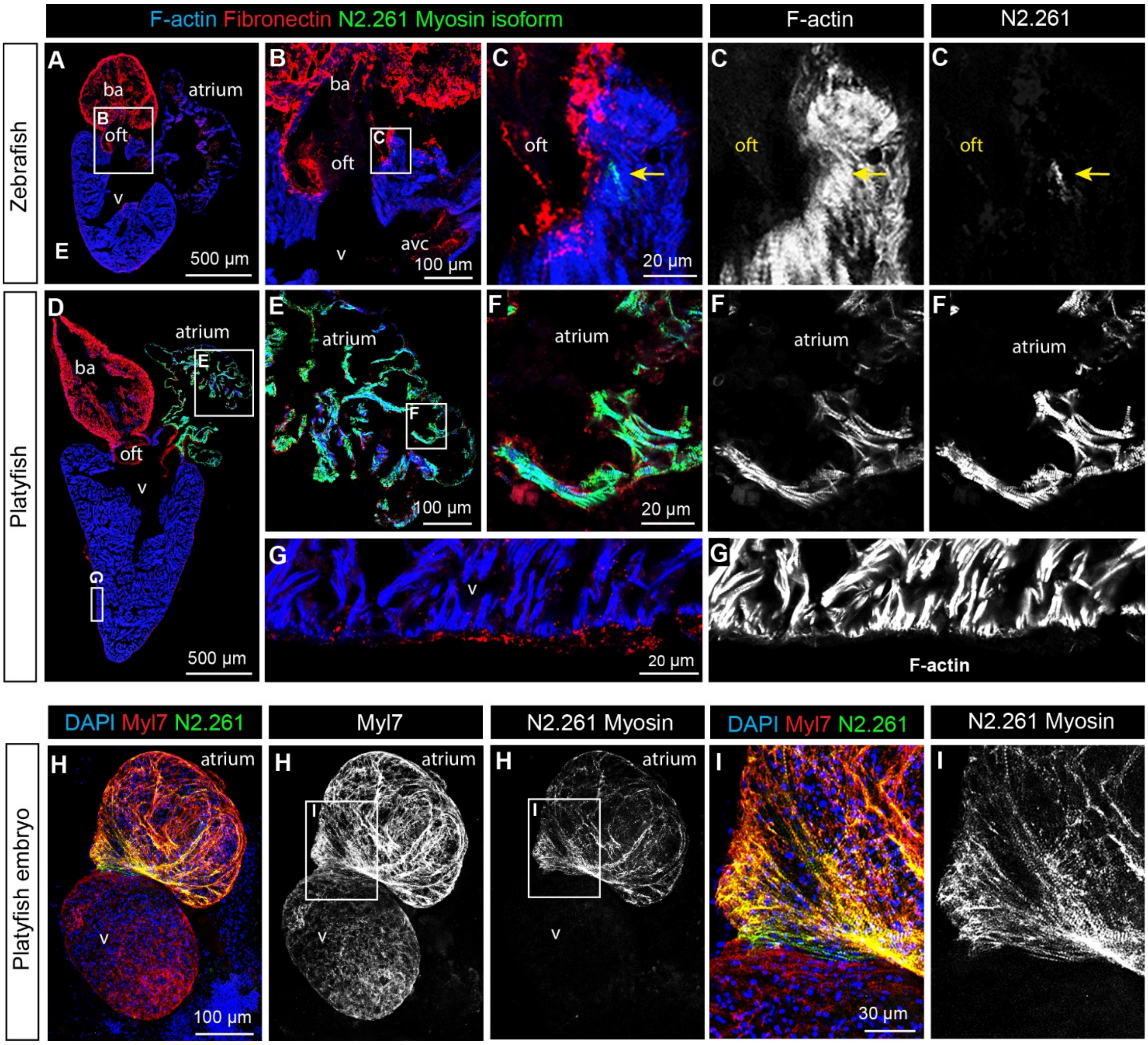


**Suppl. Figure S2. Comparison of heart morphology between zebrafish and platyfish.**
**(A–G)** Fluorescence staining of the sections shown in Figure 1C and 1F. White frames indicate regions magnified in adjacent panels, labeled with the corresponding letter. In zebrafish (B, C), N2.261 immunoreactivity is restricted to a small subset of cardiomyocytes at the outflow tract (oft). In platyfish (E, F), the N2.261 antibody immunolabels the atrium. (G) Higher magnification of the platyfish ventricular wall confirms the absence of a compact myocardial layer. avc, atrioventricular canal; ba, bulbus arteriosus; oft, outflow tract; v, ventricle.
(H–I) Whole-mount immunostaining of platyfish embryonic hearts reveals N2.261 immunoreactivity confined to the atrium.

### Supplementary Figure S3

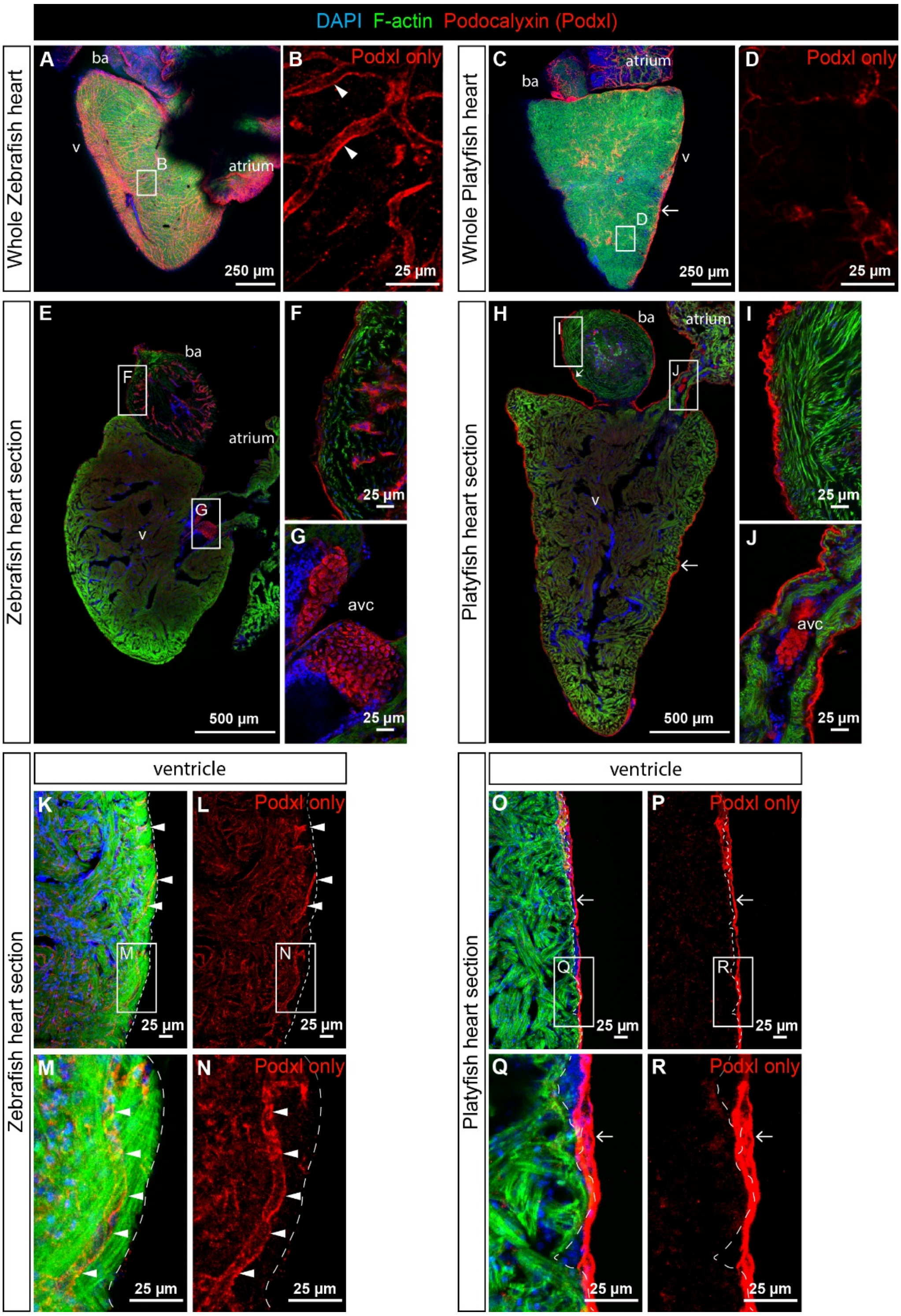


**Suppl. Figure S3. Unlike zebrafish, the platyfish ventricle lacks a vascularized myocardium.** Podocalyxin (Podxl) immunostaining, which marks the apical surface of endothelial cells lining blood vessels and the endocardium, reveals coronary vasculature in zebrafish but not in platyfish. The myocardium is labeled with F-actin; all nuclei are counterstained with DAPI.

**(A–D)** Whole-mount immunostaining of zebrafish and platyfish hearts. In zebrafish, Podocalyxin labels a dense network of blood vessels within the ventricular wall (B), which is absent in platyfish (D).

**(E–R)** Immunostaining of longitudinal heart sections from zebrafish and platyfish. Whole-heart overviews (E, H) are shown with white frames indicating regions magnified in adjacent panels,

labeled with the corresponding letters. In zebrafish, Podocalyxin marks blood vessels within the ventricular wall (K–N), the bulbus arteriosus (ba, F), and cells of the atrioventricular canal (avc, G). In platyfish, Podocalyxin is detected on the ventricular surface (O–R), the bulbus arteriosus (I), and within the valves (J); notably, no intramyocardial vascular structures are observed.
Arrowheads indicate vascular structures within the compact myocardium in zebrafish. Arrows indicate the Podocalyxin-positive ventricular surface layer in platyfish. Dashed lines demarcate the boundary of the F-actin-positive myocardium. avc, atrioventricular canal; ba, bulbus arteriosus; v, ventricle.

### Supplementary Figure S4

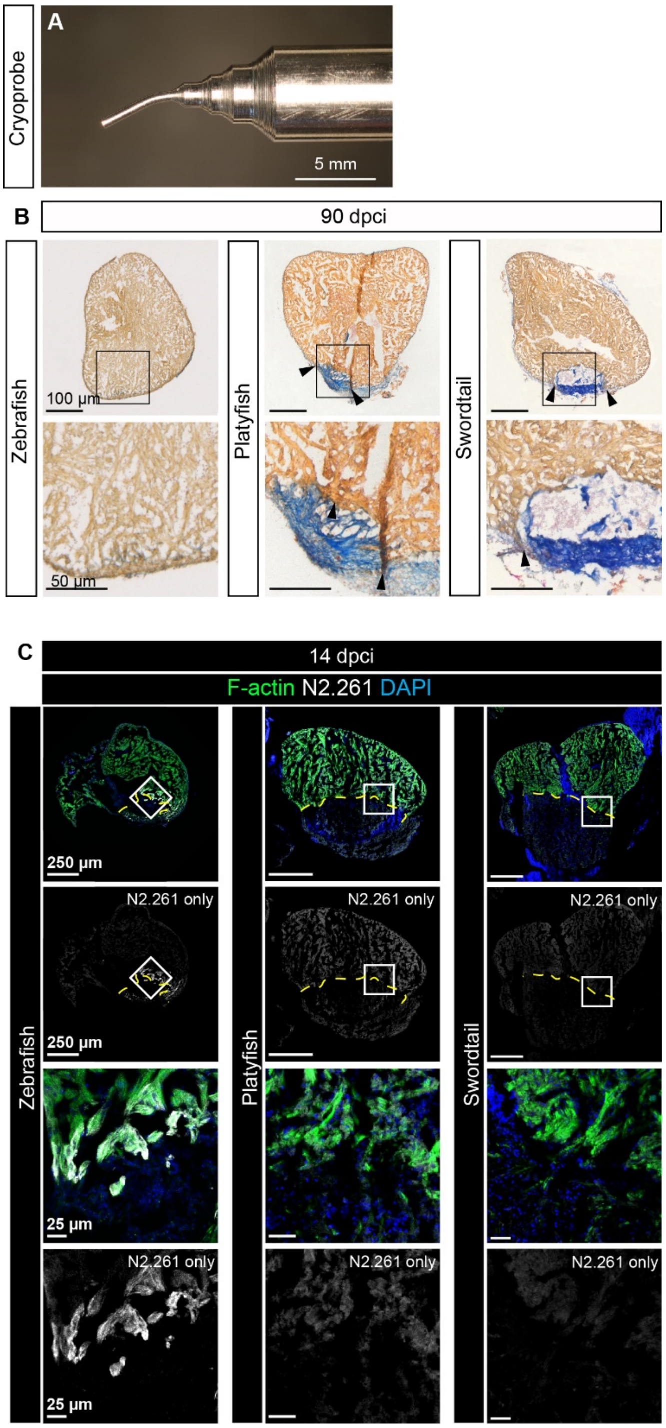


**Suppl. Figure S4. Imperfect regeneration in *Xiphophorus* fish at 90 dpci.**
**(A)** Photograph of the cryoprobe used for cardiac cryoinjury.
**(B)** AFOG staining of transverse ventricular sections from cryoinjured zebrafish (left), platyfish (middle), and swordtail (right) at 90 dpci. Intact myocardium appears orange; fibrin and protein deposits appear red; collagen appears blue. Note the persistent scar tissue covering the wound in both *Xiphophorus* species, in contrast to the regenerated zebrafish myocardium.
**(C)** N2.261 immunostaining on transverse heart sections from cryoinjured zebrafish (left), platyfish (middle), and swordtail (right) at 14 dpci. N2.261 labels dedifferentiated cardiomyocytes located approximately 100 µm from the injury site (border zone) in zebrafish, whereas no such labeling is observed in platyfish or swordtail.

**Supplementary Figure S5**

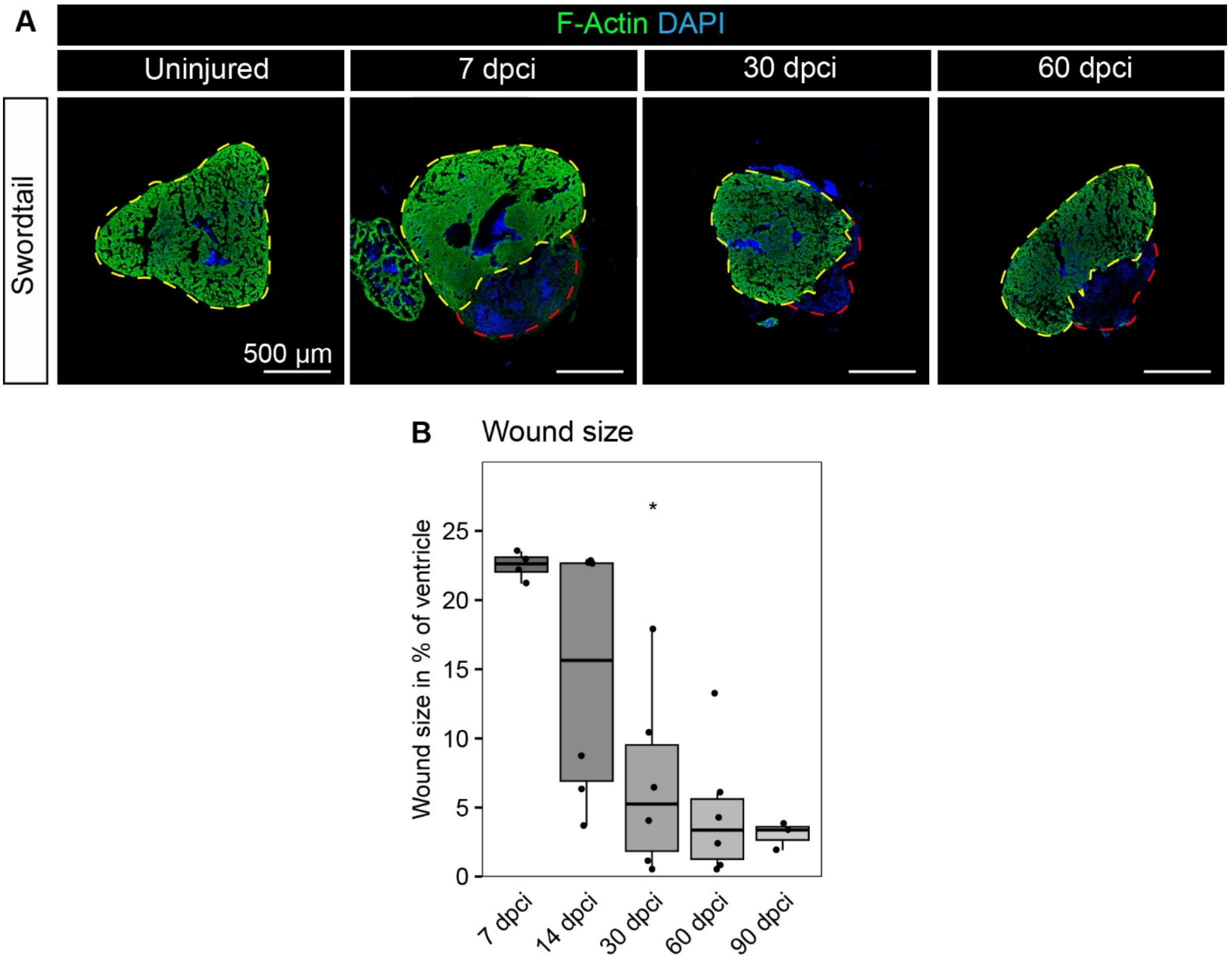


**Supplementary Figure S5: Dynamics of ventricular repair in swordtails.**
**(A)** Phalloidin-488 and DAPI staining of uninjured and cryoinjured swordtail ventricles at 7, 30, and 60 dpci. The damaged area is identified by a weak phalloidin signal reflecting the loss of contractile cardiomyocytes. Yellow dashed line surrounds the myocardium, whereas the red dashed line indicates the injury margin.
**(B)** Wound size expressed as a percentage of total ventricular volume at 7, 14, 30, 60, and 90 dpci. Statistical comparisons across time points were performed using Kruskal-Wallis tests. Adjusted p-values: *, $p < 0.05$; **, $p < 0.01$. Sample sizes: $n = 4$ (7 dpci), 6 (14 dpci), 6 (30 dpci), 6 (60 dpci), and 3 (90 dpci).

**Supplementary Figure S6**

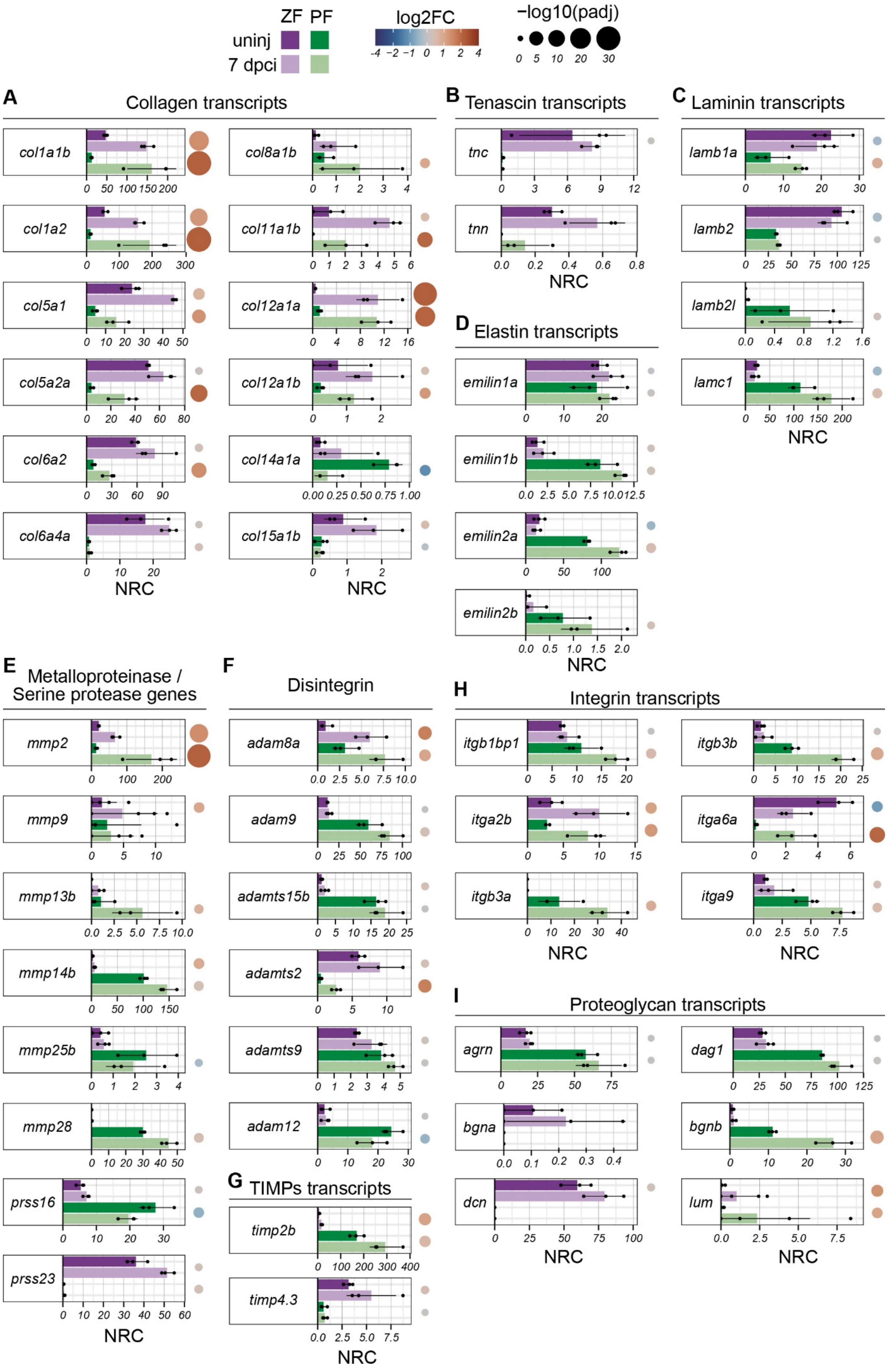

**Suppl. Figure S6: Transcript abundance of ECM-related genes in zebrafish and platyfish reveals both conserved and divergent responses to cryoinjury.**

Transcript abundance of orthologous genes was compared between zebrafish (purple) and platyfish (green) and between uninjured (dark shade) and cryoinjured (light shade) conditions for the following gene categories: (A) collagen, (B) tenascin, (C) laminin, (D) elastin, (E) metalloproteinase and serine protease, (F) disintegrin (ADAM/ADAMTS), (G) tissue inhibitor of metalloproteinase (TIMP), (H) integrin, and (I) proteoglycan genes. For details on data normalization and visualization, see the legend to Figure 5 and Methods.

### Supplementary Figure S7

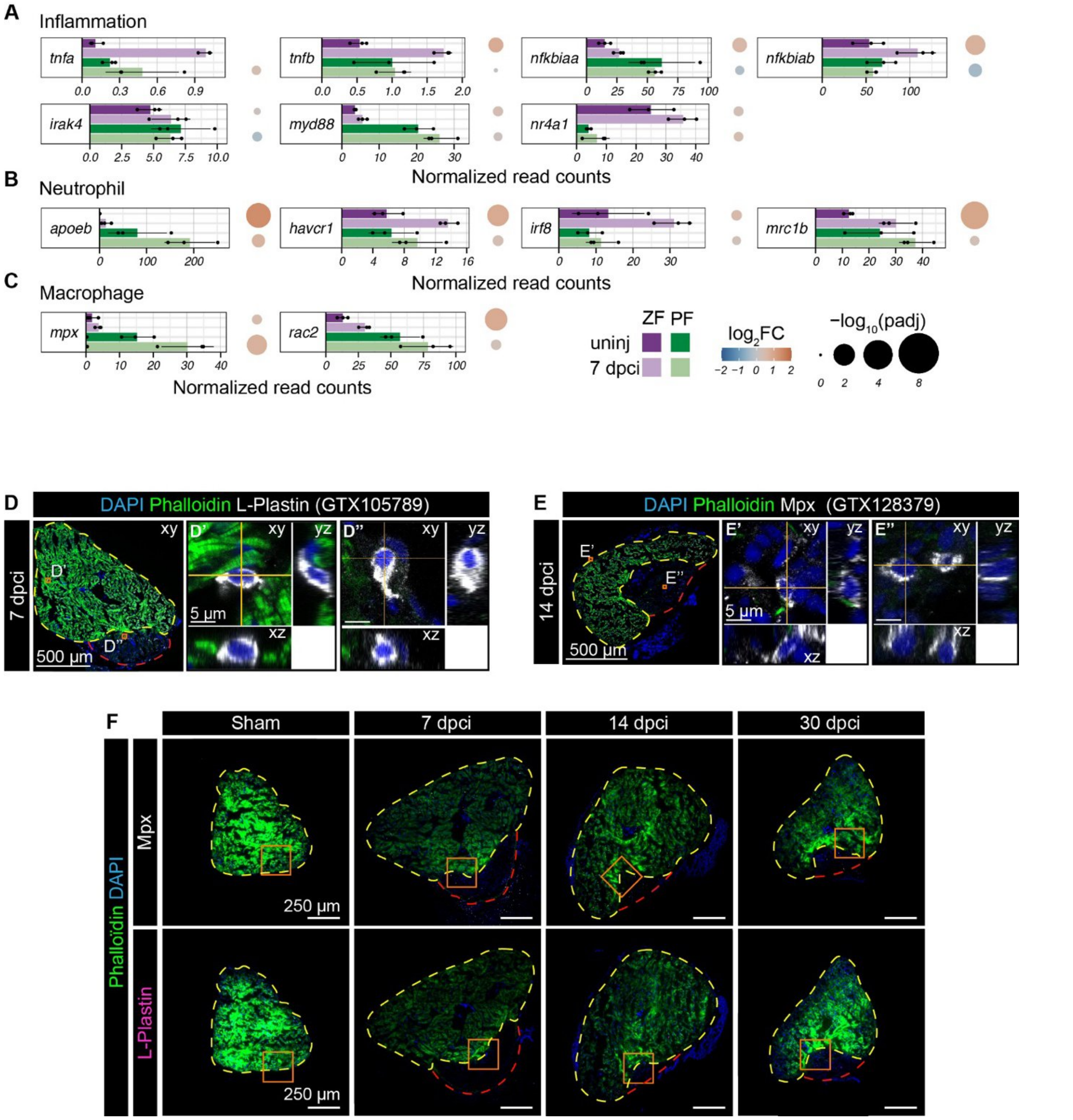


**Suppl. Figure S7. Immune cell activation and marker validation in platyfish.**
**(A–C)** Transcript abundance of genes related to inflammation (A), neutrophil recruitment (B), and macrophage recruitment (C) in zebrafish (purple) and platyfish (green) before and after cryoinjury (7 dpci). For details on data normalization and visualization, see the legend to Figure 5 and Methods.
**(D–E)** Validation of antibody specificity for L-plastin (GTX105789, D) and Mpx (GTX128379, E) immunostaining using confocal imaging. For each image, orthogonal projections are shown in the x–y, x–z, and y–z planes, confirming discrete cytoplasmic signals in muscle-negative cells.
**(F)** Widefield fluorescence images of ventricular sections from sham-operated and cryoinjured platyfish (7, 14, and 30 dpci), stained for Mpx (top row) or L-plastin (bottom row). Orange frames indicate the regions shown at higher magnification in Figure 6.
For panels (D), (E), and (F), sections were counterstained with DAPI (nuclei) and phalloidin-488 (myocardium). Dashed yellow lines demarcate the intact myocardium; dashed red lines outlines the wound area.

**Supplementary Figure S8**

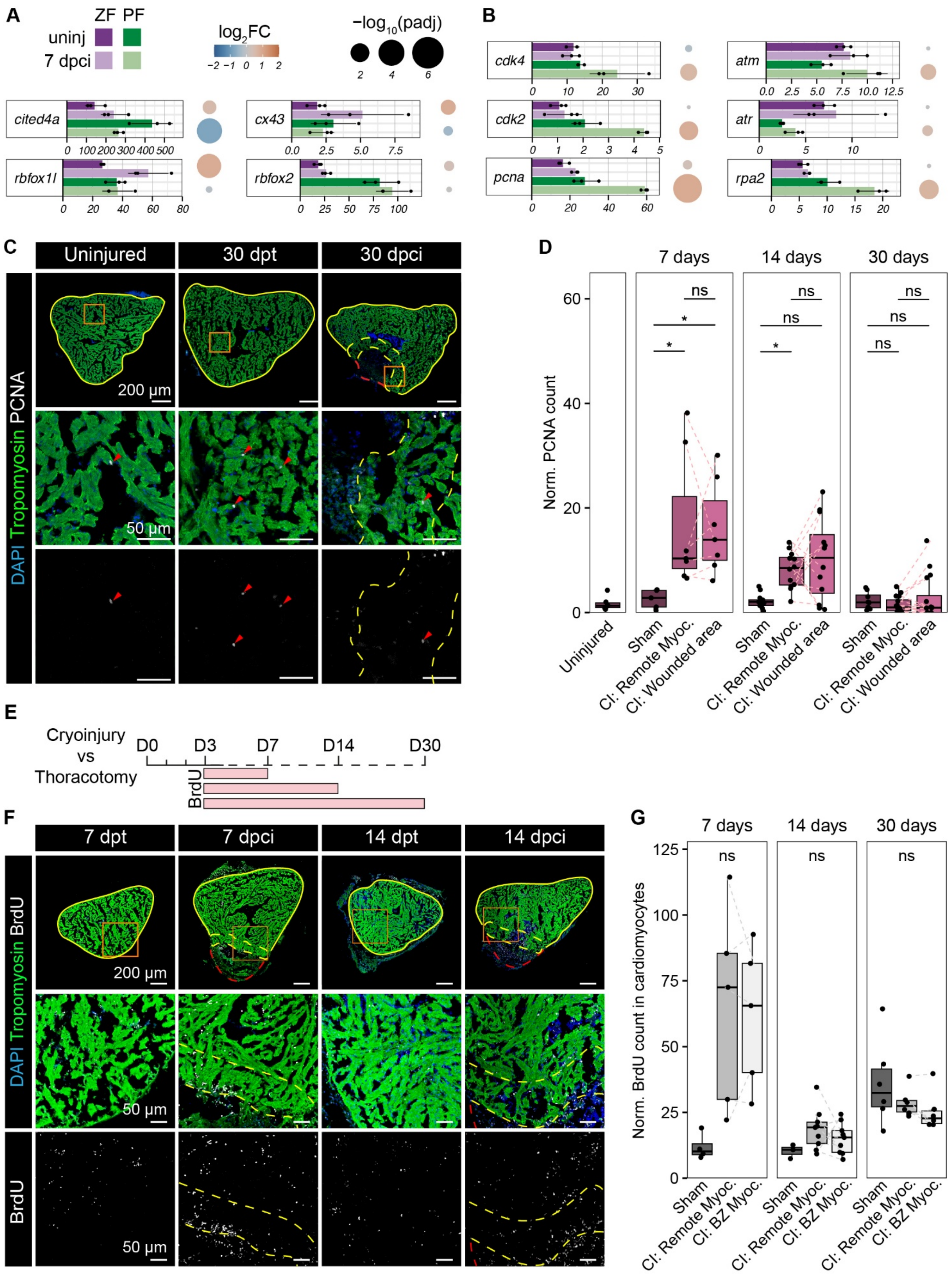

**Suppl. Figure S8: Cardiomyocytes can enter the cell cycle in platyfish.**
**(A–B)** Transcript abundance of genes related to cardiomyocyte plasticity (A) and cell proliferation (B) in zebrafish (purple) and platyfish (green) before and after cryoinjury (7 dpci). For details on data normalization and visualization, see the legend to Figure 5 and Methods.
**(C)** PCNA immunostaining in uninjured, sham-operated, and cryoinjured platyfish ventricles at 30 days post-surgery. Sections were counterstained with DAPI (nuclei) and anti-tropomyosin antibody (myocardium) to delineate the intact and wounded areas (see also panel F).
**(D)** Quantification of PCNA-positive nuclei in uninjured, sham-operated, and cryoinjured ventricles at 7, 14, and 30 days post-surgery. For cryoinjured ventricles, quantifications were performed separately in the remote myocardium (CI: Remote Myoc.) and wound area (CI: Wound). Normalized values represent the ratio of PCNA-positive nuclei to total DAPI-positive nuclei. Sample sizes: n = 6 (Uninjured), 5 (7 day - sham), 7 (7 day - CI), 9 (14 day - sham), 12 (14 day - CI), 8 (30 day - sham), 12 (30 day - CI),
**(E)** Schematic of the BrdU labeling protocol. Sham-operated and cryoinjured platyfish received continuous BrdU treatment from day 3 post-surgery until the day of heart harvest.
**(F)** BrdU immunolocalization in sham-operated and cryoinjured platyfish ventricles at 7 and 14 days post-surgery. Sections were counterstained with DAPI and anti-tropomyosin antibody.
**(G)** Quantification of BrdU-positive cardiomyocytes (CMs) in sham-operated and cryoinjured ventricles at 7, 14, and 30 days post-surgery. Cardiomyocytes were identified as tropomyosin-positive nuclei. For cryoinjured ventricles, quantifications were performed separately in the remote myocardium and border zone (CI: BZ Myoc.). Sample sizes: n = 4 (7 day - sham), 5 (7 day - CI), 3 (14 day - sham), 9 (14 day - CI), 6 (30 day - sham), 6 (30 day - CI),
For graphs in (D) and (G), Kruskal-Wallis tests were followed by pairwise Wilcoxon tests: unpaired tests were used to compare sham-operated and cryoinjured ventricles, and paired tests were used to compare different regions within the same cryoinjured heart. Holm's post hoc correction was applied for multiple comparisons.

### Supplementary Materials

#### Reagents

| Product/Molecules | Reference | Company |
|---|---|---|
| Acetic Acid Glacial | A10400 | Thermo Fisher Scientific |
| Acid fuchsin | F8129 | Sigma-Aldrich |
| Agarose | A9539 | Sigma-Aldrich |
| Aniline blue | B8563 | Sigma-Aldrich |
| BCIP | 11383221001 | Sigma-Aldrich |
| Bouin's fixative | 57211 | Thermo Fisher Scientific |
| BrdU | ab6326 | Abcam |
| DMSO | D4540 | Sigma-Aldrich |
| Entellan | 1079600500 | Merck Millipore |
| Ethanol | E10650DF/17 | Thermo Fisher Scientific |
| Formaldehyde | 252549 | Sigma-Aldrich |
| Goat serum | 005000121 | Jackson ImmunoResearch Labs |
| Heparin | H3393 | Sigma-Aldrich |
| Lidocaine hydrochloride | PAR1257 | Supelco |
| MeOH | M/400/15 | Sigma-Aldrich |
| Molybdatophosphoric Acid Hydrate (Phosphomolybdic acid) | 1.00532.0100 | Merck Millipore |
| NaCl | 207790010 | Thermo Fisher Scientific |
| NBT (Nitro Blue Tetrazolium) | 11383213001 | Sigma-Aldrich |
| TrueSeq Stranded mRNA | / | Illumina |
| Orange G | O1625 | Sigma-Aldrich |
| Phalloidin-CruzFluor-488 | sc-363791 | Santa Cruz Biotech |
| Phosphomolybdic acid | 79560 | Merck |
| Picric acid solution | P6744 | Sigma-Aldrich |
| RNase-free Dnase | | Qiagen |
| RNeasy Plus Micro Kit | | Qiagen |
| Sirius Red F3B | 3655548 | Sigma-Aldrich |
| SMART-Seq Low Input RNA Kit | | Takara Bio |
| Sodium citrate | 71402 | Fluka |
| Stainless steel cryoprobe | Home-made | |
| Steel bead | 69989 | Qiagen |
| Sucrose | SO389 | Sigma-Aldrich |
| Superfrost Plus slides | J1800AMNZ | Thermo Fisher Scientific |
| Tissue-Tek O.C.T. | 4583 | Sakura |
| Tricaine | E10521 | Sigma-Aldrich |
| Tris | T1503 | Sigma-Aldrich |
| Triton X-100 | X-100 | Sigma-Aldrich |
| TRIzol | T9424 | Sigma-Aldrich |
| Tween 20 | T2700 | Sigma-Aldrich |
| Xylene | 95662 | Sigma-Aldrich |

### Antibodies & Fluorochromes

*Primary antibodies*

| Cat# | Company | Target | Isotype & Host | Dilution | Channel used | RRID |
|---|---|---|---|---|---|---|
| A4.1025 | DSHB | MYH1 | IgG2a - Mouse | 1/100 | 488 | AB_528356 |
| ab6326 | Abcam | BrdU | IgG2a - Rat | | Cy5 | AB_305426 |
| CH1 | gift of J. Jung Chin | Tropomyosin (TPM1) | IgG1 - Mouse | 1/200 | 488 | AB_2205770 |
| F3648 | Sigma-Aldrich | Fibronectin (FN1) | Polyclonal - Rabbit | 1/400 | Cy3 | AB_476976 |
| GTX105789 | GeneTex | L-Plastin | Polyclonal - Rabbit | 1/500 | Cy5 | AB_10721688 |
| GTX128346 | Genetex | Myl7 | Polyclonal - Rabbit | 1/500 | 488 | AB_2885759 |
| GTX128379 | GeneTex | Mpx | Polyclonal - Rabbit | 1/500 | Cy5 | AB_2885768 |
| M0879 | Dako | PCNA | IgG2aK - Mouse | 1/500 | Cy5 | AB_2160651 |
| N2.261 | DSHB | embCMHC | IgG1K - Mouse | 1/50 | Cy3 | AB_531790 |
| Podocalyxin 2 | gift from M. Affolter's lab Herwig et al. 2011. Current Biology | PODXL2 | Polyclonal - Rabbit | 1/200 | Cy3 | NA |

*Secondary antibodies (All from Jackson ImmunoResearch Labs)*

| Cat# | Target | Host | Dilution | Conjugate | RRID |
|---|---|---|---|---|---|
| 112-165-167 | Rat IgG | Goat | 1/200 | Cy3 | AB_2338251 |
| 115-545-205 | Mouse IgG1 | Goat | 1/200 | Alexa Fluor 488 | AB_2338916 |
| 115-605-206 | Mouse IgG2a | Goat | 1/200 | Alexa Fluor 647 | AB_2338917 |
| 711-545-152 | Rabbit IgG | Donkey | 1/200 | Alexa Fluor 488 | AB_2313584 |
| 711-605-152 | Rabbit IgG | Donkey | 1/200 | Alexa Fluor 647 | AB_2492288 |
| 712-495-153 | Rat IgG | Donkey | 1/200 | DyLight 649 | AB_2560938 |
| 715-166-151 | Mouse IgG | Donkey | 1/200 | Cy3 | AB_2340817 |
| 715-545-151 | Mouse IgG | Donkey | 1/200 | Alexa Fluor 488 | AB_2341099 |

*Fluorochromes*

| Cat# | Company | Molecule |
|---|---|---|
| D9564 | Sigma-Aldrich | DAPI |
| sc-363791 | Santa Cruz BioTech | Phalloidin |

### Computer programs

| Software / Package | Version | Source |
|---|---|---|
| Adobe Photoshop | 27.5 | Adobe |
| Adobe Illustrator | 27.5 | Adobe |
| ClusterProfiler | 4.7.1 | Wu et al. 2021 |
| DESeq2 | 1.38.3 | Love et al. 2014 |
| dplyr | 1.1.4 | Wickham et al. 2019 |
| fastqc | 0.11.9 | Andrews 2022 |
| FeatureCounts | 2.0.1 | Liao et al. 2014 |
| Fiji/ImageJ | 1.54f | Schindelin et al. 2019 |
| forcats | 1.0.0 | Wickham et al. 2019 |
| ggnewscale | 0.5.0 | Campitelli 2024 |
| ggplot2 | 3.5.1 | Wickham et al. 2019 |
| HiSat2 | 2.2.1 | Kim et al. 2015 |
| LAS X Navigator | Different versions for different microscopes | Leica |
| Perplexity AI (assistant) | 2025 | Perplexity |
| Photoshop | 24.5.0 | Adobe |
| purrr | 1.0.4 | Wickham et al. 2019 |
| R | 4.2.1, 4.3.0 | R Core Team |
| RColorBrewer | 1.1-3 | Neuwirth 2022 |
| readr | 2.1.5 | Wickham et al. 2019 |
| RSeQC | 4.0.0 | Wang et al. 2012 |
| rstatix | 0.7.2 | Kassambara 2023 |
| Shiny | 1.6.0 | Chang et al. 2022 |
| stringr | 1.5.1 | Wickham et al. 2019 |
| tidyr | 1.3.1 | Wickham et al. 2019 |

### References for software and bioinformatics

Alboukadel Kassambara. 2023. rstatix: Pipe-Friendly Framework for Basic Statistical Tests. https://CRAN.R-project.org/package=rstatix

Andrews, S. 2022. "A Quality Control Tool for High Throughput Sequence Data." http://www.bioinformatics.babraham.ac.uk/projects/fastqc/.

Chang, Winston, Joe Cheng, J. J. Allaire, Carson Sievert, Barret Schloerke, Yihui Xie, Jeff Allen, Jonathan McPherson, Alan Dipert, and Barbara Borges. 2022. *Shiny: Web Application Framework for R*. https://CRAN.R-project.org/package=shiny.

Elio Campitelli. 2024. ggnewscale: Multiple Fill and Colour Scales in 'ggplot2'. https://CRAN.R-project.org/package=ggnewscale

Erich Neuwirth. 2022. RColorBrewer: ColorBrewer Palettes. https://CRAN.R-project.org/package=RColorBrewer

Hu, B., Lelek, S., Spanjaard, B. *et al.* Origin and function of activated fibroblast states during zebrafish heart regeneration. *Nat Genet* **54**, 1227–1237 (2022). https://doi.org/10.1038/s41588-022-01129-5

Jiang M, Xiao Y, E W, Ma L, Wang J, Chen H, Gao C, Liao Y, Guo Q, Peng J, Han X, Guo G. Characterization of the Zebrafish Cell Landscape at Single-Cell Resolution. Front Cell Dev Biol. 2021 Oct 1;9:743421. doi: 10.3389/fcell.2021.743421. PMID: 34660600; PMCID: PMC8517238.

Kanehisa, Minoru, Yoko Sato, and Masayuki Kawashima. 2022. "KEGG Mapping Tools for Uncovering Hidden Features in Biological Data." *Protein Science* 31 (1): 47–53. https://doi.org/10.1002/pro.4172.

Kim, Daehwan, Ben Langmead, and Steven L Salzberg. 2015. "HISAT: A Fast Spliced Aligner with Low Memory Requirements." *Nature Methods* 12 (4): 357–60. https://doi.org/10.1038/nmeth.3317.

Liao, Y., G. K. Smyth, and W. Shi. 2014. "featureCounts: An Efficient General Purpose Program for Assigning Sequence Reads to Genomic Features." *Bioinformatics* 30 (7): 923–30. https://doi.org/10.1093/bioinformatics/btt656.

Liberzon, Arthur, Chet Birger, Helga Thorvaldsdóttir, Mahmoud Ghandi, Jill P. Mesirov, and Pablo Tamayo. 2015. "The Molecular Signatures Database Hallmark Gene Set Collection." *Cell Systems* 1 (6): 417–25. https://doi.org/10.1016/j.cels.2015.12.004.

Love, Michael I, Wolfgang Huber, and Simon Anders. 2014. "Moderated Estimation of Fold Change and Dispersion for RNA-Seq Data with DESeq2." *Genome Biology* 15 (12): 550. https://doi.org/10.1186/s13059-014-0550-8.

R Core Team. 2022. *R: A Language and Environment for Statistical Computing*. Vienna, Austria: R Foundation for Statistical Computing. https://www.R-project.org/.

Schindelin, J., Arganda-Carreras, I., Frise, E., Kaynig, V., Longair, M., Pietzsch, T., … Cardona, A. (2012). Fiji: an open-source platform for biological-image analysis. *Nature Methods*, *9*(7), 676–682. doi:10.1038/nmeth.2019

Subramanian, Aravind, Pablo Tamayo, Vamsi K. Mootha, Sayan Mukherjee, Benjamin L. Ebert, Michael A. Gillette, Amanda Paulovich, et al. 2005. "Gene Set Enrichment Analysis: A Knowledge-Based Approach for

Interpreting Genome-Wide Expression Profiles." *Proceedings of the National Academy of Sciences* 102 (43): 15545–50. https://doi.org/10.1073/pnas.0506580102.

Wang, Liguo, Shengqin Wang, and Wei Li. 2012. "RSeQC: Quality Control of RNA-Seq Experiments." *Bioinformatics* 28 (16): 2184–5. https://doi.org/10.1093/bioinformatics/bts356.

Wickham H, Averick M, Bryan J, Chang W, McGowan LD, François R, Grolemund G, Hayes A, Henry L, Hester J, Kuhn M, Pedersen TL, Miller E, Bache SM, Müller K, Ooms J, Robinson D, Seidel DP, Spinu V, Takahashi K, Vaughan D, Wilke C, Woo K, Yutani H (2019). "Welcome to the tidyverse." *Journal of Open Source Software*, **4**(43), 1686. doi:10.21105/joss.01686.

Wu, Tianzhi, Erqiang Hu, Shuangbin Xu, Meijun Chen, Pingfan Guo, Zehan Dai, Tingze Feng, et al. 2021. "clusterProfiler 4.0: A Universal Enrichment Tool for Interpreting Omics Data." *The Innovation* 2 (3): 100141. https://doi.org/10.1016/j.xinn.2021.100141.

Zhou Q, Zhao C, Yang Z, Qu R, Li Y, Fan Y, Tang J, Xie T, Wen Z. Cross-organ single-cell transcriptome profiling reveals macrophage and dendritic cell heterogeneity in zebrafish. Cell Rep. 2023 Jul 25;42(7):112793. doi: 10.1016/j.celrep.2023.112793. Epub 2023 Jul 14. PMID: 37453064.
